# Comparative developmental transcriptomics of *Drosophila* mushroom body neurons highlights the mevalonate pathway as a regulator of axon growth

**DOI:** 10.1101/2025.09.17.676757

**Authors:** Lora Fahdan, Hagar Meltzer, Noa Wigoda, Ron Rotkopf, Oren Schuldiner

## Abstract

The ability of neurons to extend axons is governed by tightly regulated genetic programs that vary across developmental stages and cell types. Understanding the molecular features that control axon growth potential is critical for uncovering how neural circuits form, mature, and respond to injury or disease. The *Drosophila* mushroom body (MB) offers a powerful model to dissect axon growth programs, as lineage-related Kenyon cells (KCs) undergo different developmental events under shared spatiotemporal conditions. During metamorphosis, γ-KCs undergo axon pruning, followed by developmental regrowth at the same time-frame as α/β-KCs initiate axon growth - thus providing a unique opportunity to compare these distinct growth paradigms. We thus performed RNA-sequencing of α/β-and γ-KCs during their initial growth and developmental regrowth, respectively, revealing dynamic transcriptional changes and identifying 300 shared genes upregulated during both growth states. A targeted loss-of-function screen revealed genes specifically required for either α/β initial growth, γ regrowth, or both. Focusing on one such candidate, Pmvk, we found that it plays a crucial role in axon regrowth by acting within the mevalonate pathway. Notably, other enzymes in this pathway were also required, suggesting that the entire metabolic pathway is essential for supporting regrowth. Genetic mutant analyses and rescue exepriements suggest that Pmvk likely controls axon regrowth via Rheb, an effector of the TOR pathway, which we previously found to be required for regrowth. Our developmental transcriptomic atlas not only advances understanding of intrinsic axon growth programs, but also provides candidate genes and a valuable framework for future studies aimed at enhancing axon regeneration in the adult nervous system.

## Introduction

Axon growth is essential for correct circuit formation during development. It is a tightly controlled process, and an emerging paradigm claims that axon growth and synaptogenesis are mutually exclusive processe (Bradke, 2022; Hilton et al., 2022). Despite its significance, the cellular and molecular mechanisms that govern different axon growth states are not fully understood. Axon growth capacity is also crucial during regeneration following injury. Santiago Ramon y Cajal (Rozo et al., 2024) first noted that while the adult mammalian central nervous system typically fails to regenerate, the peripheral nervous system can sprout and grow new connections. Regeneration is influenced by both intrinsic factors that determine axon growth potential, and extrinsic factors such as deposits by glia or other cells (Sutherland and Geoffroy, 2020; Zheng and Tuszynski, 2023). Therefore, understanding how intrinsic axon growth capacity is mechanistically regulated in different developmental contexts should also forward our understanding of what limits axon regeneration in pathological states such as spinal cord injury.

*Drosophila* is an excellent model organism to investigate axon growth potential in different developmental contexts, due to its unparalleled neurogenetic toolkit combined with the extensive remodeling of its nervous system during metamorphosis. Neuronal remodeling is a conserved strategy to refine neural circuits, by eliminating exuberant connections and promoting regrowth toward adult-specific targets (Schuldiner and Yaron, 2015). The *Drosophila* mushroom body (MB), a center of associative learning and memory in the fly brain (Belle and Heisenberg, 1994), undergoes stereotypic remodeling during early pupal stages. The MB is comprised of three major types of intrinsic neurons: the γ-, α′/β′-and α/β-Kenyon Cells (KCs), sequentially born from four neuroblasts per hemisphere (Lee et al., 1999). The first-born γ-KCs initially extend axons that bifurcate to form the larval vertical and medial lobes. However, in early pupae (initiated at ∼6 hours after puparium formation; h APF), both axonal lobes are pruned back to their branch point, followed by axon regrowth (initiated at ∼24h APF) to form the adult-specific γ lobe that only projects medially (Lee et al., 1999). In contrast, the later-born α′/β′-and α/β-KCs retain their original bifurcated projections into adulthood (Figure 1A). Importantly, the initial growth of α/β-axons occurs at ∼24h APF – the same time in which γ-axons initiate developmental regrowth.

**Figure 1.**
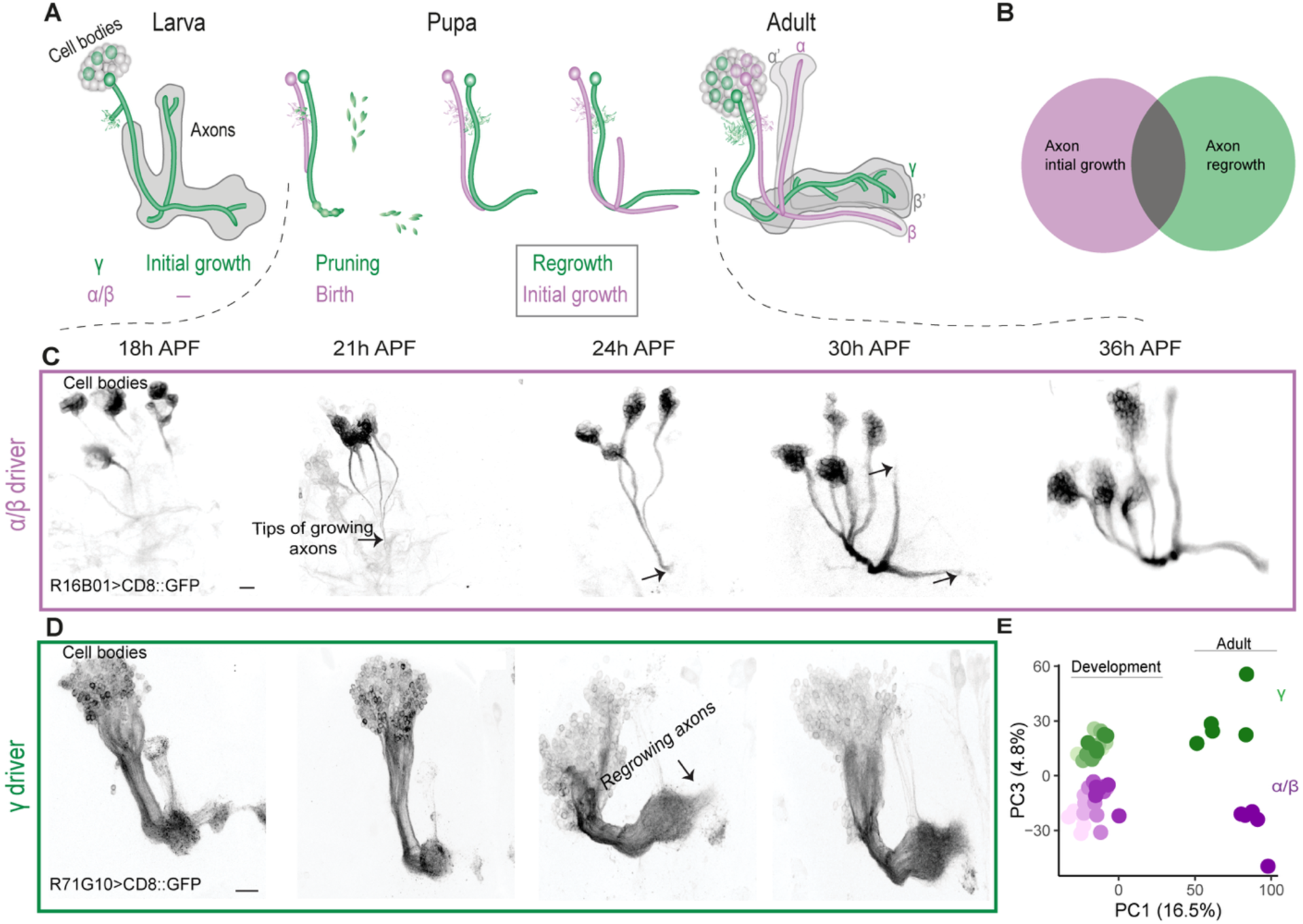
Lineage-related neuronal types undergo intial axon growth and regrowth in shared spatiotemporal conditions. (A) A model of MB development highlighting that the regrowth of γ-axons temporally and spatially overlaps with the initial growth of α/β-axons. (B) A Venn diagram depicting the theoretical and predicted comparison between α/β initial axon growth and γ-axon regrowth. (C) Confocal Z-projections of R16B01-Gal4 driving expression of membranal GFP (mCD8::GFP), which specifically labels developing α/β-KCs during the pupal stage. The images were captured at five time points between 18h until 36h APF, when the axons reach the end of the lobes. At 18h APF, four groups of cell bodies can be observed, corresponding to newly born cells from the four neuroblasts, each sending out short axons that progressively extend in each subsequent image. The axon tips are marked with arrows. (D) Confocal Z-projections of γ-KCs labeled with R71G10-Gal4 driving mCD8::GFP, as identified by (Alyagor et al., 2018), between 18h to 30h APF. At 21h APF, the axons are completely pruned and begin to re-extend for a second time. By 24h APF, the regrowing axons are visible (arrow). (E) PCA of α/β and γ-KC expression profiles during development and in adult. Color code depicts neuronal type (green for γ and magenta for α/β) and developmental stage (from light to dark as development progresses).

**Figure 2.**
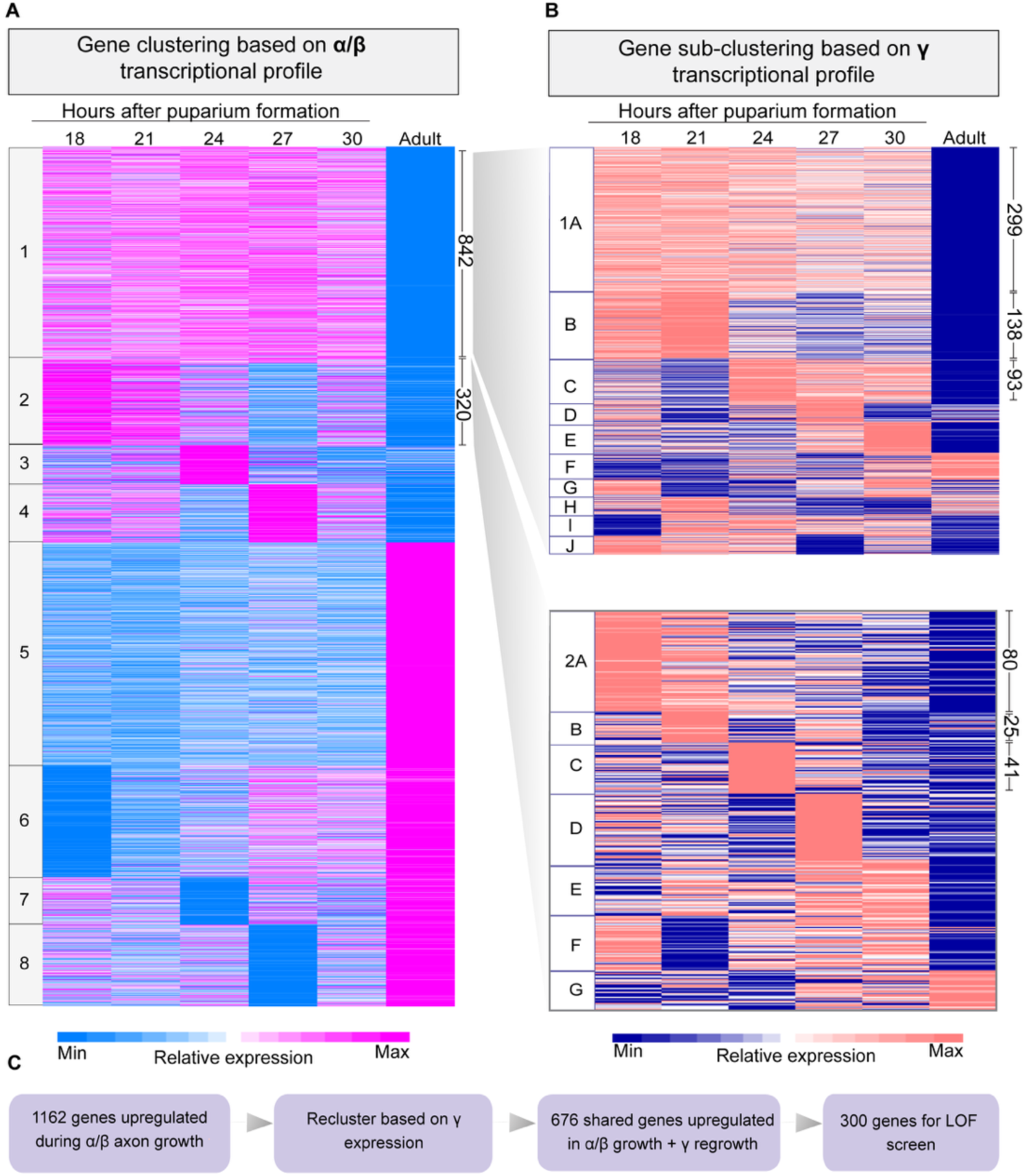
Comparative transcriptomic analysis of α/β-axon initial growth and γ-axon regrowth. (A) Heatmap showing k-means clustering of 3345 differentially expressed genes during α/β-KC development. Light blue and magenta represent low and high relative expression, respectively. (B) Heatmaps showing k-means re-clustering of 842 genes from cluster 1 or 320 genes from cluster 2 based on their γ-KC transcriptional profiling. Blue and coral represent low and high relative expression, respectively. For all the heatmaps, each horizontal line describes the relative expression of a single gene. Gene expression was determined in five timepoints after pupa formation (18, 21, 24, 27 and 30 hours APF) and in adult. The number of genes within selected clusters are specified to the right. A flow chart detailing the criteria used to select candidate genes for the loss-of-function (LOF) screen.

**Figure 3.**
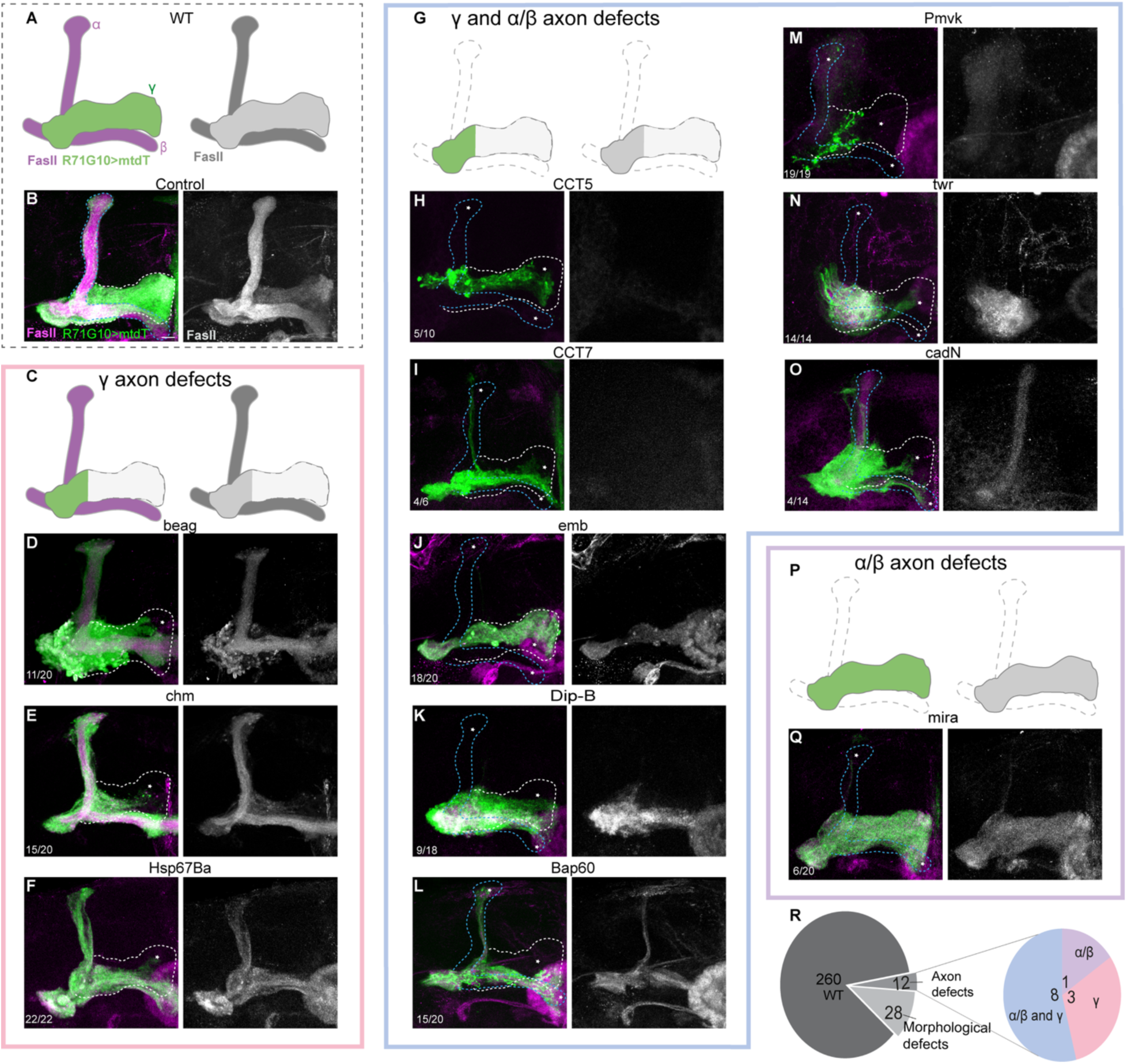
RNAi-based screen identifies 12 genes that are essential for γ-and/or α/β-axon morphology. (A) Schematic of the WT α/β-and γ-axonal lobes, illustrating labeling patterns that match the corresponding confocal images. γ-axons are labeled in green using mtdT:3xHA under the control of R71G10-QF2. FasII is visualized with antibody staining shown in magenta, and also presented separately in greyscale. (B) Confocal Z-projection of a control MB in which γ-KCs are labeled by mtdT:3xHA (green) driven by R71G10-QF2, and OK107-Gal4 drives the expression of Luciferase. Magenta (left panel) or greyscale (right panel) is anti-FasII staining, which strongly labels α/β-axons and weakly labels γ-axons. (C,G,P) Schematics of α/β-and/or γ-axon lobe defects. (D-F,H-O,Q) Confocal Z-projections of adult MBs in which γ-KCs are labeled by mtdT:3xHA (green) driven by R71G10-QF2, with OK107-Gal4 driving expression of RNAi targeting beag (beag; D); chameau (chm; E); Heat shock gene 67Ba (Hsp67Ba; F); Chaperonin containing TCP1 subunit 5 (CCT5; H); Chaperonin containing TCP1 subunit 7 (CCT7; I); embargoed (emb; J); Dipeptidase B (Dip-B; K); Brahma associated protein 60kD (Bap60; L); Phosphomevalonate kinase (Pmvk; M); twisted bristles roughened eye (twr; N); Cadherin-N (cadN; O); miranda (mira; Q). (R) Pie charts showing the phenotypic distribution of the 300 screened genes (left), and of the axon growth phenotypes of the 12 ‘positive hits’ (right). The putative WT γ-and α/β-lobes are outlined in white and blue, respectively. Asterisks mark the missing part of α/β or γ lobes. Frame colors in (C-Q) represent the phenotypic group as depicted in (R). Numbers represent the fraction of MBs displaying the presented phenotype. Scale bar is 10 μm.

γ-axons thus undergo dynamic transitions between intial growth, pruning, and regrowth, all of which were shown to be tightly regulated by nuclear receptor complexes (Lee et al., 2000; Rabinovich et al., 2016; Yaniv et al., 2012), implying they are goverened by genetic programs. We previously employed RNA-sequencing to characterize the transcriptional landscape of γ-KCs throughout development at high temporal resolution (Alyagor et al., 2018). Moreover, our lab demonstrated that the molecular mechanisms underyling developmental regrowth are distinct from those of initial axon growth, yet show similarities to regeneration after injury (Marmor-Kollet and Schuldiner, 2016; Rabinovich et al., 2016; Sudarsanam et al., 2020; Yaniv et al., 2020). The fact that different axon growth states share core machinery yet also show distinct pathways depending on the cellular and developmental context, motivated us to directly compare the genetic programs of initial axon growth and developmental axon regrowth.

In this study, we took advantage of the temporal overlap between α/β initial growth and γ regrowth to compare the transciptional landscape of these distinct axon growth programs occuring in a similar developmental, environmental and lineage context (Figure 1A-B). This comparison laid the groundwork for a large-scale loss-of-function screen aimed at uncovering genes and pathways required for both axon growth paradigms. Through this approach, we identified 12 genes that are required for initial axon growth and/or regrowth. We delved deeper into phosphomevalonate kinase (Pmvk), an enzyme in the mevalonate pathway, and found that it likely promotes axon regrowth, but not initial growth, at least in part via the TOR pathway. Together, our findings reveal shared and divergent molecular pathways underlying distinct axon growth programs, offering new insights into how intrinsic factors shape neuronal growth capacity. By combining time-course transcriptional profiling of defined neuronal populations with functional genetics, we not only uncovered key regulators of axon regrowth, but also provide a rich, publicly available dataset that can serve as a foundation for future discoveries in neuronal development.

## Methods

### Key resources table

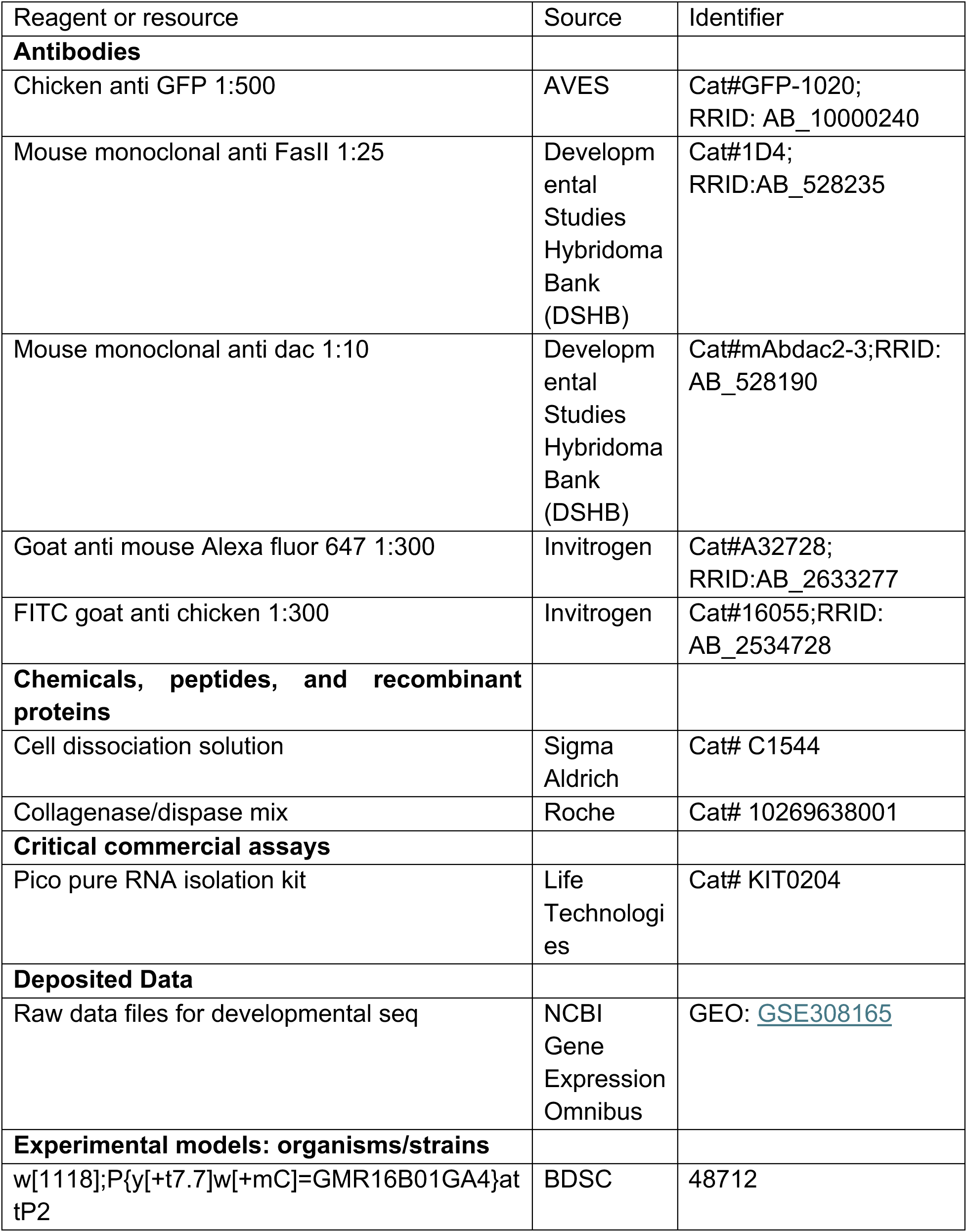
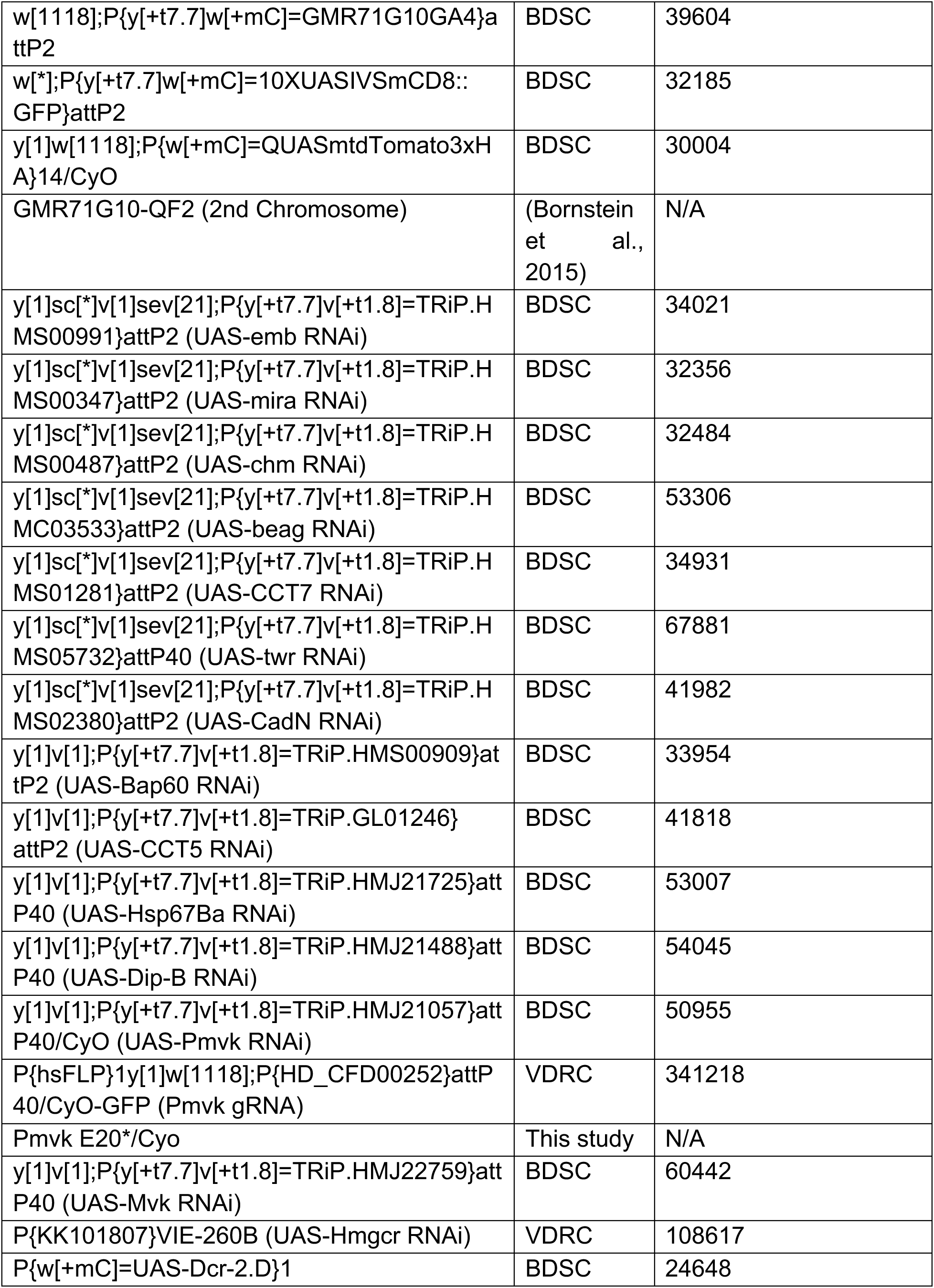
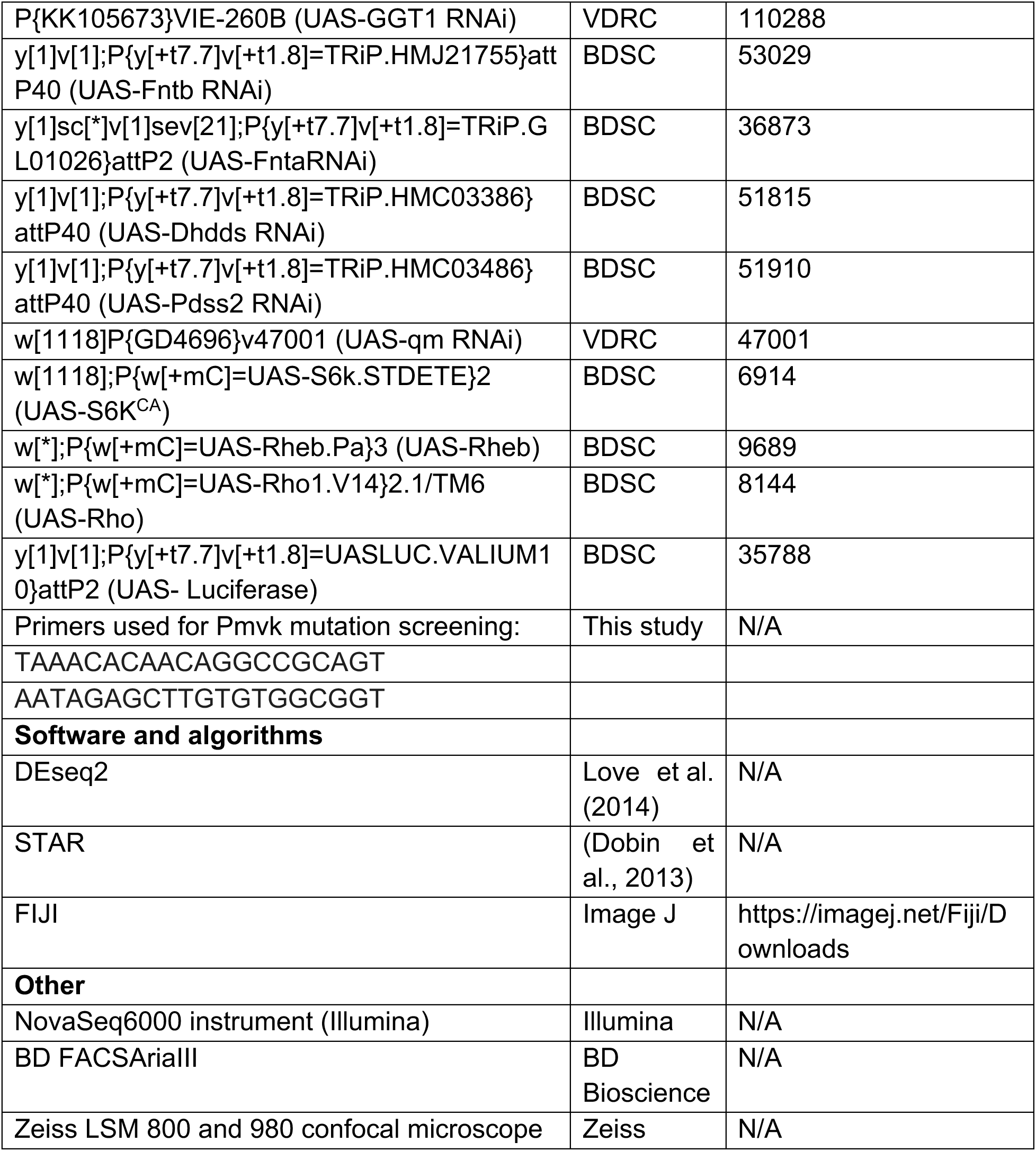

### Fly husbandry and strains

All fly strains were reared at 25°C (unless indicated otherwise) under standard laboratory conditions on molasses-containing food. Unless specifically written, similar numbers of male and female flies were taken. ‘Adult’ refers to flies 3-5 days post eclosion.

The 30 driver lines screened to specifically label α/β-KCs are described in Supplementary Table 1. The RNAi lines used in the loss-of-function screen are detailed in Supplementry Table 3.

RNAi lines were obtained from the Vienna Drosophila Resource Center (VDRC), Austria, or from Bloomington Drosophila Stock Center (BDSC), USA.

### Genotypes

hsFLP is y,w,hsFLP122; 40A and 82B are FRTs on 2L and 3R respectively; 71G10 is GMR71G10-GAL4; 71G10-QF2 is GMR71G10-QF2; OK107 is OK107-GAL4; R44E04 is GMR44E04-GAL4; 16B01 is GMR16B01-Gal4; G80 is TubP-Gal80. QUAS-mtdT is QUAS-mtdt:HA, UAS-CD8:GFP is UAS-mCD8::GFP.

Males and females were used interchangeably.

Figure1:

(I) (C) UAS-CD8:GFP/+;71G10/+

(II) (D) UAS-CD8:GFP/+;16B01/+

Figure3:

(B) 71G10-QF2,QUAS-mtdT/+;UAS-Luciferase/+;OK107/+

(D) 71G10-QF2,QUAS-mtdT/+;UAS-beag RNAi^TRiP.HMC03533^/+;OK107/+

(E) 71G10-QF2,QUAS-mtdT/+;UAS-chm RNAi^TRiP.HMS00487^/+;OK107/+

(F) 71G10-QF2,QUAS-mtdT/+;UAS-Hsp67Ba RNAi^TRiP.HMJ21725^/+;OK107/+

(H) 71G10-QF2,QUAS-mtdT/+;UAS-CCT5 RNAi^HMC02988^/+;OK107/+

(I) 71G10-QF2,QUAS-mtdT/+;UAS-CCT7 RNAi^TRiP.HMS01281^/+;OK107/+

(J) 71G10-QF2,QUAS-mtdT/+;UAS-emb RNAi^TRiP.GL01246^/+;OK107/+

(K) 71G10-QF2,QUAS-mtdT/+;UAS-Dip-B RNAi^TRiP.HMJ21488^/+;OK107/+

(L) 71G10-QF2,QUAS-mtdT/+;UAS-Bap60 RNAi^TRiP.HMS00909^/+;OK107/+

(M) 71G10-QF2,QUAS-mtdT/+;UAS-Pmvk RNAi^TRiP.HMJ21057^/+;OK107/+

(N) 71G10-QF2,QUAS-mtdT/+;UAS-twr RNAi^TRiP.HMS05732^/+;OK107/+

(O) 71G10-QF2,QUAS-mtdT/+;UAS-cadN RNAi^TRiP.HMS02380^/+;OK107/+

(Q) 71G10-QF2,QUAS-mtdT/+;UAS-mira RNAi^TRiP.HMS00347^/+;OK107/+

Figure 4:

(A,C) 71G10,UAS-CD8:GFP/+;UAS-Luciferase/+

(B,D) 71G10,UAS-CD8:GFP/+;UAS-Pmvk RNAi ^TRiP.HMJ21057^/+

(E) R44E04,UAS-CD8:GFP/+;UAS-Luciferase/+

(F) R44E04,UAS-CD8:GFP/+;UAS-Pmvk RNAi ^TRiP.HMJ21057^/+

(G,I) hsFLP,UAS-CD8::GFP/+;G80,40A/40A;71G10/+

(H,J) hsFLP,CD8:GFP/+;UAS-PmvkRNAi ^TRiP.HMJ21057^(attP40)/+;82B,Gal80/82B;OK107/+

(K) hsFLP,UAS-CD8::GFP/+;G80,40A/40A;R44E04/+

(L) hsFLP,UAS-CD8::GFP/+;G80,40A/*Pmvk^E20*^*,40A;R44E04/+

Figure 5:

(A) 71G10-QF2,QUAS-mtdT/+;UAS-Luciferase/+;OK107/+

(B) UAS-Dicer;71G10-QF2,QUAS-mtdT/+;UAS-Hmgcr RNAi KK 101807/+;OK107/+

(C) 71G10-QF2,QUAS-mtdT/+;UAS-Mvk RNAi ^TRiP.HMJ22759^/+;OK107/+

Figure 6:

(C) hsFLP,UAS-CD8::GFP/+;G80,40A/40A;71G10/+

(D) hsFLP,UAS-CD8::GFP/+;G80,40A/*PmvkE20*\*,40A;71G10/+

(C) (E) hsFLP,UAS-CD8::GFP/+;G80,40A/*PmvkE20*\*,40A;71G10/UAS-Rho

(D) (F) hsFLP,UAS-CD8::GFP/+;G80,40A/*PmvkE20*\*,40A;71G10/UAS-Rheb

(E) (G) hsFLP,UAS-CD8::GFP/+;G80,40A/*PmvkE20*\*,40A;71G10/UAS-S6KCA

Figures S3:

(A) hsFLP,UAS-CD8:GFP;G80,40A/40A;OK107/+

(B) hsFLP,UAS-CD8:GFP;G80,40A/40A;UAS-emb RNAi^TRiP.HMS00991^/+;OK107/+

(C) hsFLP,UAS-CD8:GFP;G80,40A/40A;UAS-CCT5 RNAi^TRiP.GL01246^/+;OK107/+

(D) hsFLP,UAS-CD8:GFP;G80,40A/40A;UAS-CCT7 RNAi^TRiP.HMS01281^/+;OK107/+

(E) hsFLP,UAS-CD8:GFP;G80,40A/40A;UAS-CadN RNAi^TRiP.HMS02380^/+;OK107/+

(F) hsFLP,CD8:GFP/+;UAS-Pmvk RNAi ^TRiP.HMJ21057^/+;82B,G80/82B;OK107/+

(G) hsFLP,CD8:GFP/+;UAS-twr RNAi^TRiP.HMS05732^/+;82B,G80/82B;OK107/+

(H) hsFLP,UAS-CD8:GFP;G80,40A/40A;UAS-Bap60 RNAi^TRiP.HMS00909^/+;OK107/+

(I) hsFLP,CD8:GFP/+;UAS-Dip-B RNAi^TRiP.HMJ21488^ /+;82B,G80/82B;OK107/+

(J) hsFLP,UAS-CD8:GFP;G80,40A/40A;UAS-beag RNAi^TRiP.HMC03533^/+;OK107/+

(K) hsFLP,UAS-CD8:GFP;G80,40A/40A;UAS-chm RNAi^TRiP.HMS00487^/+;OK107/+

(L) hsFLP,CD8:GFP/+;UAS-Hsp67Ba RNAi^TRiP.HMJ21725^ /+;82B,G80/82B;OK107/+

(M) hsFLP,UAS-CD8:GFP;G80,40A/40A;UAS-mira RNAi^TRiP.HMS00347^/+;OK107/+

Figures S4:

(A) hsFLP,UAS-CD8:GFP;G80,40A/40A;OK107/+

(B) hsFLP,UAS-CD8:GFP;G80,40A/40A;UAS-emb RNAi^TRiP.HMS00991^/+;OK107/+

(C) hsFLP,UAS-CD8:GFP;G80,40A/40A;UAS-CCT5 RNAi^TRiP.GL01246^/+;OK107/+

(D) hsFLP,UAS-CD8:GFP;G80,40A/40A;UAS-CCT7 RNAi^TRiP.HMS01281^/+;OK107/+

Figure S5:

(B) 71G10-QF2, QUAS-mtdT/+;UAS-Luciferase/+;OK107/+

(B) UAS-Dicer;71G10-QF2,QUAS-mtdT/+;UAS-GGT1 RNAi KK 105673/+;OK107/+ (D)71G10-QF2,QUAS-mtdT/+;UAS-Fntb RNAi ^TRiP.HMJ21755^/+;OK107/+

(E) 71G10-QF2,QUAS-mtdT/+;UAS-Fnta RNAi ^TRiP.GL01026^/+;OK107/+

(F) 71G10-QF2,QUAS-mtdT/+;UAS-Dhdds RNAi ^TRiP.HMC03386^/+;OK107/+

(G) 71G10-QF2,QUAS-mtdT/+;UAS-Pdss2 RNAi ^TRiP.HMC03486^/+;OK107/+

(H) 71G10-QF2,QUAS-mtdT/+;UAS-qm RNAi GD 4696/+;OK107/+

Figure S6:

(A) hsFLP,UAS-CD8::GFP/+;G80,40A/40A;71G10/+

(B) hsFLP,UAS-CD8::GFP/+;G80,40A/40A;71G10/UAS-Rheb

(C) hsFLP,UAS-CD8::GFP/+;G80,40A/40A;71G10/UAS-Rho

(D) hsFLP,UAS-CD8::GFP/+;G80,40A/40A;71G10/UAS-S6KCA

Figure S7:

(A) hsFLP,UAS-CD8::GFP/+;G80,40A/40A;71G10/+

(B) hsFLP,UAS-CD8::GFP/+;G80,40A/*Pmvk^E20*^*,40A;71G10/+

(C) hsFLP,UAS-CD8::GFP/+;G80,40A/*Pmvk^E20*^*,40A;71G10/UAS-Rho

(D) hsFLP,UAS-CD8::GFP/+;G80,40A/*Pmvk^E20*^*,40A;71G10/UAS-Rheb

(E) hsFLP,UAS-CD8::GFP/+;G80,40A/*Pmvk^E20*^*,40A;71G10/UAS-S6K^CA^

## Materials and methods

### Cell dissociation and sorting

Brains were dissected in a cold Ringer’s solution and incubated at 29°C with a collagenase/dispase mix for 15 minutes for larval and pupal brains, and 30 minutes for adult brains. After enzymatic digestion, the tissue was washed with a dissociation solution (Sigma-Aldrich) and mechanically dissociated into single cells. The cell suspension was then passed through a 35μm mesh (Falcon) to remove debris and cell clusters. A total of 1000 GFP+ cells were sorted using a BD FACSAriaIII (BD Biosciences) equipped with a 100μm nozzle under low pressure, directly into 100μl of Pico-Pure RNA isolation kit extraction buffer (Life Technologies). RNA extraction was subsequently performed using the kit, or frozen at-80°C for future use. To minimize potential injury responses, samples were kept on ice throughout the procedure, from dissection to RNA extraction except from the dissociation enzyme incubation.

### RNA amplification and library preparation Bulk smart-seq2

Bulk RNA-seq libraries were prepared at the Crown Genomics Institute of the Nancy and Stephen Grand Israel National Center for Personalized Medicine (G-INCPM), Weizmann Institute of Science. Libraries were prepared based on the smart-seq2 protocol (Picelli et al., 2014) with small modifications. Briefly, 250 pg of RNA was used as input. Reverse transcription was performed using Oligo-dT primers, followed by PCR (16 cycles). After Agencourt Ampure XP beads cleanup (Beckman Coulter), the amplified cDNA underwent library generation using Nextera XT (Illumina). Libraries were quantified by Qubit (Thermo Fisher Scientific) and TapeStation (Agilent). Sequencing was done on a NovaSeq6000 instrument (Illumina) using an SP 100 cycles kit, allocating approximately 13M reads per sample (single read sequencing).

### Analysis of RNA-Seq data

Two sequencing runs of RNA were combined, yielding a mean of 22.5 million reads (single-read sequencing) and a minimum of 10 million reads per sample. The reads were aligned with STAR **<u>(DOI: 10.1093/bioinformatics/bts635)</u>** v2.5.2 to the reference genome of *Drosophila melanogaster* (DM6, UCSC). Gene annotations were obtained from FlyBase.org (Dmel r6.46). Gene expression levels were quantified using STAR. 12,493 genes had been identified in total across the two datasets (in either γ or α/β or both). Out of these, only genes that were expressed in at least four samples (replicates) of a single time point (condition) were kept, so that the 8,221 genes that met this requirement were used in further analysis. To get relevant p-values and fold-changes for comparisons of a condition with all-zeros vs. a condition with non-zero value, zero values were replaced with randomly assigned numbers between 1 and 7. The updated count matrix was subjected to normalization and statistical analysis using the UTAP2 pipeline (Lindner et al., 2025) which took batch effects into account and employed the DESeq2 algorithm (Love et al., 2014). Genes were considered differentially expressed if they had an adjusted p-value (padj) ≤ 0.05, a fold change ≥ 1.5 and a mean read counts of ≥10 reads.

### Clusters and sub-clusters generation and analysis

Based on the α/β transcriptome, gene expression at each of the five developmental stages (18, 21, 24, 27, and 30 hours APF) was compared to the adult stage. Genes were classified as differentially expressed during development if, at any time point, their expression was at least 1.5-fold different from the adult expression, with an adjusted p-value (padj) < 0.05 and a mean read count of at least 10, resulting in the identification of 3,345 genes. Using Partek^TM^ Genomics Suite (version 7), we performed k-means clustering on the mean expression across replicates at each developmental time point, selecting k = 8 after testing multiple values, as this produced the most distinct and coherent clusters. For subclustering the α/β Cluster 1 and Cluster 2, we employed k-means clustering based on the average expression of those genes in γ-KCs across replicates at each time point. Cluster 1 was further subdevided into 10 additional subclusters 1A-J, and cluster 2 into 7 subclusters 2A-G. We identified statistically enriched terms using Metascape (Zhou et al., 2019). This tool integrates multiple authoritative databases to identify enriched Gene Ontology (GO) terms, canonical pathways, and functional gene groups. For each cluster, the input gene lists were analyzed against the default background, and significantly enriched categories were reported based on a hypergeometric test with multiple-testing correction. The resulting enrichment networks and bar plots were used to interpret the biological processes and pathways associated with each gene cluster.

### Immunohistochemistry

Brains were dissected in cold Ringer’s solution, fixed with 4% paraformaldehyde (PFA) for 20 minutes at room temperature (RT), and subsequently washed in phosphate buffer (PB) containing 0.3% Triton-X (PBT). The washing protocol consisted of three rapid rinses followed by three additional washes, each lasting 20 minutes. To minimize non-specific staining, the samples were incubated in PBT supplemented with 5% heat-inactivated goat serum. They were then treated with primary antibodies overnight at 4°C, followed PBT washes (as above), then incubation with secondary antibodies for 2 hours at RT, followed by PBT washes. The brains were then mounted using SlowFade (Invitrogen) and visualized using Zeiss LSM800 or LSM980 confocal microscopes. Image analysis was carried out with ImageJ 1.51 (NIH).

### Generation of CRISPR mutant flies

To generate the Pmvk mutant, we used a fly strain (VDRC#341218) that expresses two guide RNAs (sgRNAs) under the control of a UAS promoter. These sgRNAs target coding exons in the 5′ region of the Pmvk gene, without overlapping the start codon. This design facilitates the introduction of small insertions or deletions (indels) in the presence of Cas9. To drive germline mutagenesis, we incorporated a nanos-Gal4 on the third chromosome and crossed these flies with a UAS-Cas9 expressing fly, which also carries a 40A FRT site on the second chromosome. After two generations, individual males were crossed with balancer flies for mutation screening. We recovered a Pmvk mutant allele that contained a nucleotide substitution (G>T) at position 60 (*Pmvk^E20*^*, Figure 4G).

**Figure 4.**
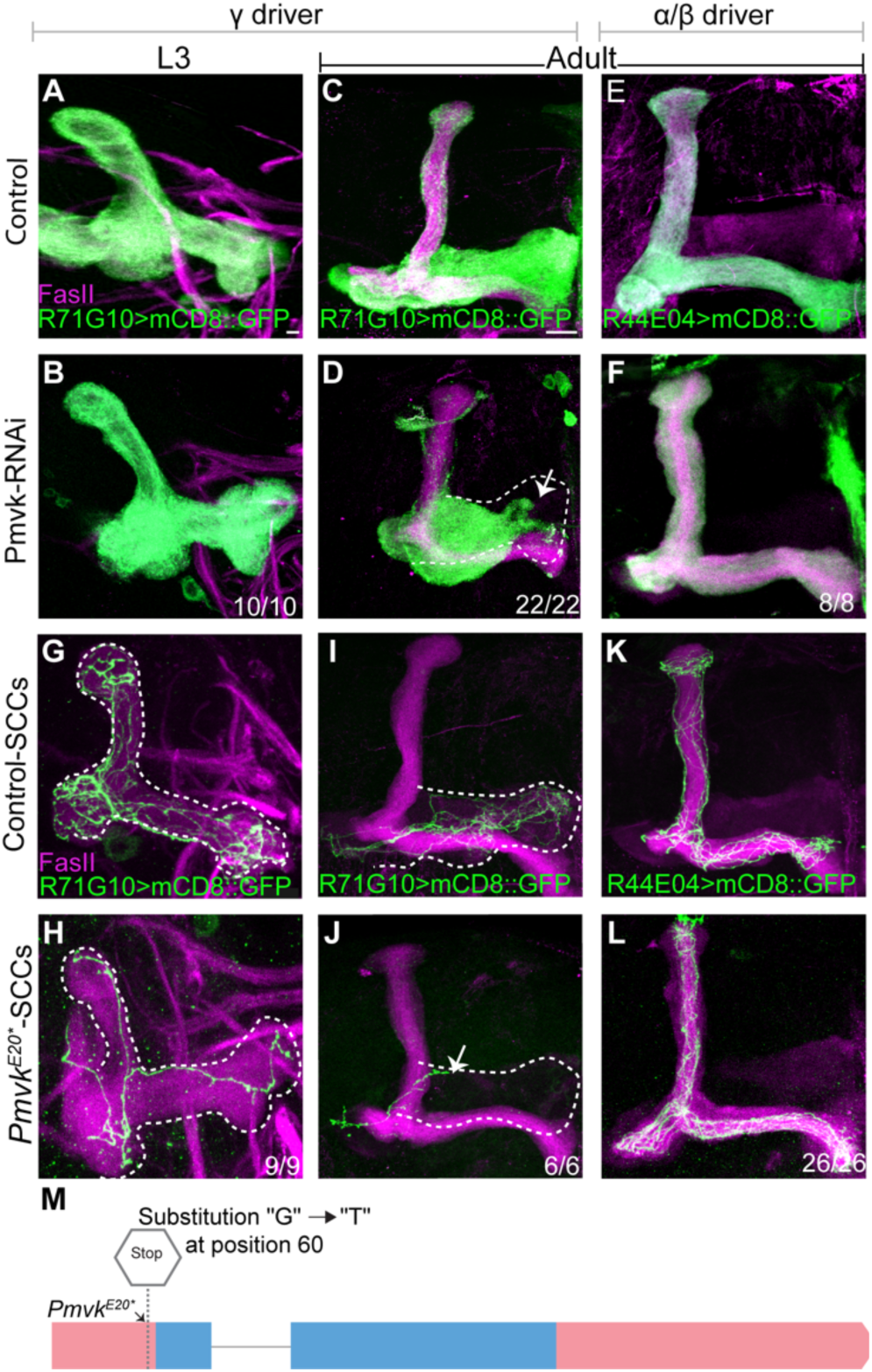
Pmvk is cell autonomously required for γ-axon regrowth but not for initial axon extension of γ nor α/β. (A-F) Confocal Z-projections of L3 (A-B) or adult (C-F) MBs with γ-KCs labeled by mCD8::GFP (green) driven by the γ-specific R71G10-Gal4 (A-D), or the α/β-specific R44E04-Gal4 (E-F). MBs in images (B-F) also express RNAi targeting Pmvk driven by R71G10-Gal4 (B,D) or R44E04-Gal4 (F). (G-L) Confocal z-projections of control (G,I,K) or *Pmvk^E20*^* (H,J,L) MARCM clones in L3 (G-H) or Adult (I-L). R71G10-Gal4 (G-J) or R44E04-Gal4 (K-L) drive expression of mCD8::GFP (green). Clones were generated by heat-shocking at 24 hours after egg-laying (G-J) or at 0h APF (K-L). The full extent of the γ lobe in adult and L3 is outlined in white (as determined by the FasII staining; magenta). Arrows demarcate axon growth defects. Scale bar is 10 μm. Numbers represent the fraction of MBs displaying the presented phenotype. (M) Schematic representation of the *Pmvk^E20*^* mutant allele, which includes a single-nucleotide substitution (G>T) at position 60, corresponding to amino acid 20, leading to the replacement of glutamic acid (E) with a stop codon.

### MARCM analysis

Mosaic Analysis with a Repressible Cell Marker (MARCM) clones were generated in a similar strategy as previously described (Lee and Luo, 1999). To generate γ-or α/β-clones, newly hatched larvae (NHL) or pupae at 0h APF, respectively, underwent a heat shock treatment at 37°C for 40 minutes. Larvae and pupae were then transferred to a 25°C incubator where they remained until eclosion.

## Data and software availability

The data discussed in this publication have been deposited in NCBI’s Gene Expression Omnibus and are accessible through GEO series accession number GEO: GSE308165 (https://www.ncbi.nlm.nih.gov/geo/query/acc.cgi?acc=GSE308165)

## Regrowth quantification

To quantify regrowth, we used a method described in a previous publication (Bornstein et al., 2021) and illustrated in Figure 6. Two reference lines were extended: line y (red), representing the full length of the γ-lobe (determined based on FasII staining), and line x (yellow), indicating the extent of clonal γ-axon regrowth (determined as the most distal point the axons reach). The regrowth index was calculated as the ratio of x to y (x/y), providing a normalized measure of axonal regrowth.

Statistical analyses for regrowth quantification were performed in R, v.4.5.1. Proportional data were logit-transformed prior to testing to meet assumptions of normality and homogeneity of variance. Group differences were evaluated using one-way ANOVA, followed by Dunnett’s post hoc test comparing all groups to Pmvk. A significant difference was detected only between control and *Pmvk^E20*^* (p < 0.001).

## Results

### Lineage-related neuronal types in the *Drosophila* mushroom body undergo intial axon growth and developmental regrowth in similar spatiotemporal conditions

In a previous study (Alyagor et al., 2018), we found that the developmental age was by far the most dominant factor in defining the transcriptional profile of γ-KCs - a principle that was later observed in developing projection neurons as well (Xie et al., 2021). Remarkably, adult γ-KCs were more similar, on a global level, to other adult neurons than to larval or pupal γ-KCs. Notably, while initial axon growth occurs in embyonic and early larval life, developmental regrowth occurs during metamorphosis. Therefore, to reliably compare the genetic programs of initial axon growth and developmental regrowth, we sought to profile two similar neuronal types that undergo different axon growth states at the same developmental window. The lineage-related α/β- and γ-KCs were a perfect match, as within a similar timeframe during the pupal stage, α/β-KCs extend their axons for the first time, while γ-KCs regrow their axons after pruning (Figure 1A-B).

To achieve our goal, we screened for drivers that would specifically label α/β-KCs at the pupal stage, when they send out axonal processes. We screened 30 Gal4 driver lines considered to be α/β-specific in adult flies (Supplementary Table 1), and imaged them at 24h APF. To our surprise, only one line - R16B01-Gal4 - was specific to the MB (i.e., not expressed elsewhere in the brain), and within the MB, specific to α/β-KCs throughout their growth (Figure 1C). At 18h APF, α/β-KCs labeled with R16B01-Gal4 showed cell bodies with short, loosely arranged axons that were still difficult to follow. By 21h APF, these axons had gathered into a tight bundle, and their growing axon tips were visible. Observations throughout the pupal stage revealed that these axons extended and reached the tips of the lobes by 36h APF (Figure 1C).

Identifying this driver enabled us to isolate α/β-KCs throughout their initial axon growth. For comparative purposes, we also repeated the isolation and profiling of γ-KCs, as previously done in our lab (Alyagor et al., 2018), using the same driver that specifically labels developing γ-KCs (R71G10-Gal4; Figure 1D). Labeling both types of neurons gave us the opportunity to isolate and sequence each population while they extend their axons for the first (α/β) or second (γ) time. Altogether, we sorted and sequenced α/β-and γ-KCs at 5 pupal timepoints-every three hours from 18h until 30h APF - in addition to adult.

We used Principal Component Analysis (PCA) to compare the transcriptional profiles of developing and adult α/β-and γ-KCs. Our results confirmed our prior knowledge (Alyagor et al., 2018) that adult γ-or α/β-KCs are more similar to each other than to their own developmental forms. Notably, developing γ-and α/β-KCs occupied a specific region in the PCA space (Figure 1E).

### Comparative transcriptomic analysis of α/β-axon initial growth and γ-axon regrowth

Our bulk smart-seq2 (see materials and methods) dataset includes reads from 8,221 distinct genes expressed in at least in one cell type (γ and/or α/β) across six developmental time points. To identify developmentally-regulated patterns of gene expression, we first analyzed the expression profiles of α/β-KCs. We thus concentrated on 3,345 genes that demonstrated dynamic expression in α/β-KCs, i.e., showed statistically significant differential expression during at least one developmental timepoint compared to adult. k-means clustering of those α/β-KC dynamic expression patterns resulted in eight distinct clusters with unique developmental expression profiles (see materials and methods). For example, genes in clusters 1 and 2 are upregulated during the pupal stage but show decreased expression in adults, and thus potentially include activators of axon growth (Figure 2A). In contrast, cluster 5 genes are downregulated during the pupal stage and upregulated in adults, suggesting that they are required for adult-specific functions. Gene Ontology (GO) term analysis further supports this observation. Cluster 1 is enriched for genes involved in axon development and other key developmental processes, including Toll pathway signaling, cytoskeletal rearrangement, and PTEN regulation. In contrast, cluster 5 is enriched for genes associated with synaptic transmission as well as learning and memory (Figure S2).

As α/β clusters 1+2 include the best candidates for axon growth promoters but contain a large number of genes (1162 genes; Figure 2A, C), we sought to narrow down the list of candidates. Since we aimed to identify genes required for axon growth in both neuronal types, we decided focus on genes within α/β clusters 1+2 and examine their expression in the γ-KC dataset. We therefore re-clustered the 1,162 genes in α/β clusters 1+2 based on their transcriptional expression in γ-KCs (Figure 2B): cluster 1 was subdivided into clusters 1A-1J, while cluster 2 was subvidided into 2A-2G. To identify shared genes required for both the initial growth of α/β-axons and the regrowth of γ-axons (Figure 2B– C), we focused on the first three sub-clusters in each heatmap (clusters 1A-C; 2A-C), as they are comprised of genes that are upregulated during axon growth of both neuronal types. From these six clusters, we compiled a set of 676 genes, out of which we selected a subset of 300 genes, which are strongly upregulated during axon growth, associated with biologically relevant functions based on GO term analysis, and have availabe tools to perturb their function (namely – flies expressing RNAi targeting these specific genes). Genes primarily associated with processes less directly related to axon growth—such as cell cycle regulation, DNA replication, chromosome organization, chromatin remodeling, and ribosomal structure—were excluded from further analysis.

### RNAi-based screen identifies 12 genes that are essential for γ-and/or α/β-axon morphology

For the loss-of-function screen, we employed an RNAi-mediated knockdown (KD) strategy, where we expressed RNAi species targeting candidate genes using the OK107-Gal4 driver (which is strongly expressed in all KC types - γ, α’/β’ and α/β). γ-axons were visualized in adult brains using a second, independent binary system, via expression of myristylated tandem Tomato (mtdT) driven by R71G10-QF2 – which consistently labels γ-KCs and stochastically labels α/β-KCs. Due to this stochasticity, we additionally stained the brains with anti-Fasciclin II (FasII) antibody, which strongly labels α/β-axons (and weakly labels γ-axons, but never α’/β’-axons).

Out of the 300 genes we screened, KD of 40 genes resulted in various morphological and structural defects (such as thinner axonal lobes, axon misguidance, and branching abnormalities; list of genes in Supplementary Table 2; phenotypic data not shown), 12 of which showed axonal defects – which we categorized into three phenotypic groups (Figure 3): defects in γ-axons but normal α/β-axons (Figure 3C-F); defects in both α/β-and γ-axons (Figure 3G-O); and normal γ-axons but no visible α/β axons (Figure 3P-Q). In addition to the axonal phenotypes that are categorized in Figure 3, KD of several of these 12 genes also resulted in additional morphological defects, such as altered number and shape of cell bodies (data not shown). Notably, 7 of the 12 genes identified in the screen are involved in protein maturation, encompassing post-translational modifications such as folding, cleavage, and prenylation.

### Pmvk, cadN and twr are cell-autonomously required for γ-axon regrowth

To inspect the KD phenotypes at better resolution, as well as gain information regarding cell-autonomous requirement, we silenced the genes specifically within neuroblast (NB) clones using the mosaic analysis with a repressible cell marker (MARCM) technique (Lee and Luo, 1999). We used the pan-KC driver OK107-Gal4, resulting in NB clones in which all KC types are labeled as well as express the relevant RNAi. To our surprise, we observed that only Pmvk, cadN and twr showed severe defects in γ-axon regrowth (Figure S3E-G), while the others exhibited γ lobes that were either WT-like, or showed weak regrowth defects (Figure S3). For mira KD, which we expected to affect α/β growth, the lobes seemed to form normally (Figure S3M). In the case of emb, CCT5, and CCT7, the neuroblast clones included only γ-, but no α/β-or α’/β’-KCs (Figure S3B-D), most likely due to a proliferation defect of the neuroblast, which prevented it from generating later born neurons.

The proliferation defects in the emb, CCT5 and CCT7 NB clones might suggest that the lack of α/β axons upon KD in the entire population of neurons (Figure 3) is because the neurons were never born, rather than a *bona fida* axon growth defect. To more specifically examine the role of these three candidate genes in α/β-axon growth, we selectively silenced emb, CCT5 and CCT7 in α/β-KCs by inducing MARCM clones at 0h APF (Figure S4) - a time in which α/β-KCs are born (Lee and Luo, 1999). Only single cell clones were visible, suggesting cell-lethality or severe proliferation defects. Importantly, all α/β single-cell clones exhibited WT-like axonal growth (Figure S4), indeed suggesting that the axonal defects observed in Figure 3 are due to non-cell-autonomous effects.

Taken together, our findings indicate that Pmvk, cadN, and twr are cell-autonomously required for γ-axon regrowth. For the purpose of this study, we chose to focus on Pmvk, although cadN and twr remain compelling candidates for future investigation.

### Pmvk is cell autonomously required for γ-axon regrowth but not for initial axon extension of γ- nor α/β-KCs

To further investigate the role of Pmvk in axon growth during development, we specifcially silenced it in γ-KCs using the γ-specific driver R71G10-Gal4. We found that while γ-axons extended normally during the larval stage (L3; Figure 4A-B), the medial γ-lobe failed to fully regrow as evident in the adult stage (Figure 4C-D). To determine whether Pmvk is also required for α/β-axon growth, we silenced it using the R44E04-Gal4 driver (of note, R44E04-Gal4 is strongly expressed in α/β throughout development but also transienty in non-MB neurons, which is why it was not chosen for the RNA profiling experiment; Supplementary Table 1). In contrast to the γ-axons, α/β-axons showed no detectable defects following Pmvk KD (Figure 4E-F).

We next used CRISPR/Cas9 to generate a novel Pmvk mutant allele - *Pmvk^E20^* - which carries a single-nucleotide substitution resulting in a premature stop codon at amino acid 20 (Figure 4M). Due to its homozygous lethality, we analyzed this mutant allele in MARCM clones. In *Pmvk^E20^* single-cell γ clones, initial axon growth appeared normal (Figure 4G- H), but, as expected, axons failed to regrow following pruning (Figure 4I-J). In contrast to γ-axons, and in accordance with the RNAi KD results, α/β neuroblast *Pmvk^E20^* clones displayed normal axon elongation (Figure 4K-L). Notably, Pmvk mutants produced no NB clones, suggesting an additional requirement for Pmvk in neuroblast proliferation or cell survival.

### The mevalonate pathway is essential for γ-axon regrowth

Pmvk is part of the mevalonate pathway – a highly conserved metabolic enzymatic pathway that consumes acetyl coenzyme A to synthesize a wide variety of isoprenoids, such as cholesterol, dolichol, and ubiquinone (Figure 5D; Edwards and Ericsson, 2025; Hinson et al., 1997). We knocked down the additional seven key enzymes in the pathway and found that pan-KC silencing of either Hmgcr or Mvk led to seemingly similar γ- regrowth defects (but not defects in ɑ/β extension; Figure 5A-C), suggesting an overall requirement of the mevalonate pathway for γ-axon regrowth.

**Figure 5.**
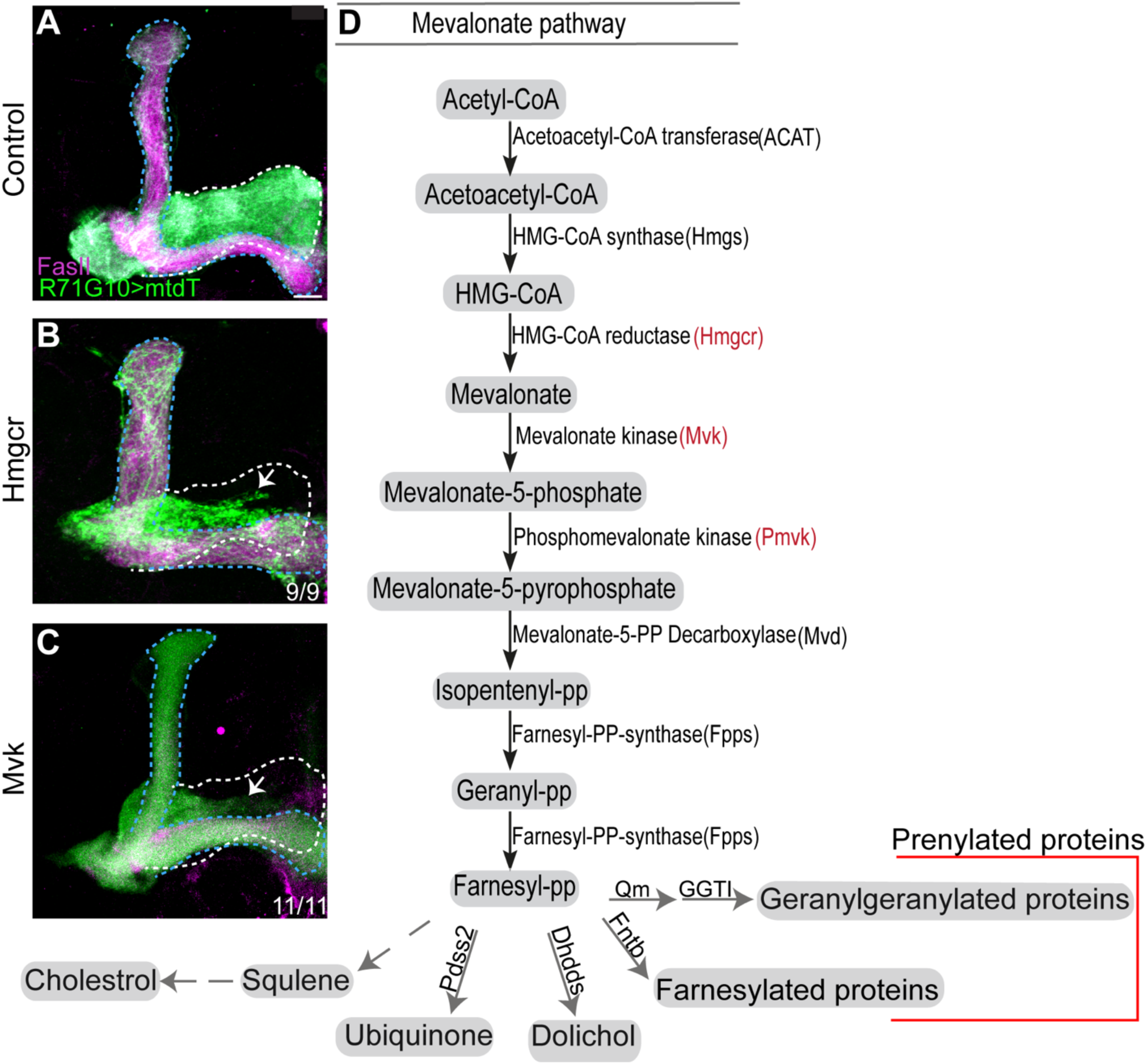
**The mevalonate pathway is essential for γ-axon regrowth**. (A-C) Confocal Z-projections of adult MBs from control (A) or those expressing RNAi targeting Hmgcr (B) or Mvk (C) driven by the pan-KC driver OK107-Gal4. γ-KCs are labeled by mtdT:3xHA (green) driven by R71G10-QF2. Magenta is FasII staining. The putative WT α/β- and putative γ-lobes are outlined in blue and white, respectively. Arrows indicate axon growth defects. Numbers represent the fraction of MBs displaying the presented phenotype. Scale bar is 10 μm. (D) A schematic of the mevalonate pathway, illustrating key metabolites and enzymes. Enzymes shown in red are those found to influence γ-axon development. The diagram also highlights the different downstream routes that FPP can take—either toward the synthesis of sterol isoprenoids (e.g., cholesterol) or non-sterol isoprenoids (such as ubiquinone, dolichol, and isopentenyl-tRNA). Additionally, FPP serves as a precursor for protein prenylation, which occurs in two main forms: farnesylation and geranylgeranylation. The dashed arrows emphasize that cholesterol biosynthesis is absent in *Drosophila*

**Figure 6.**
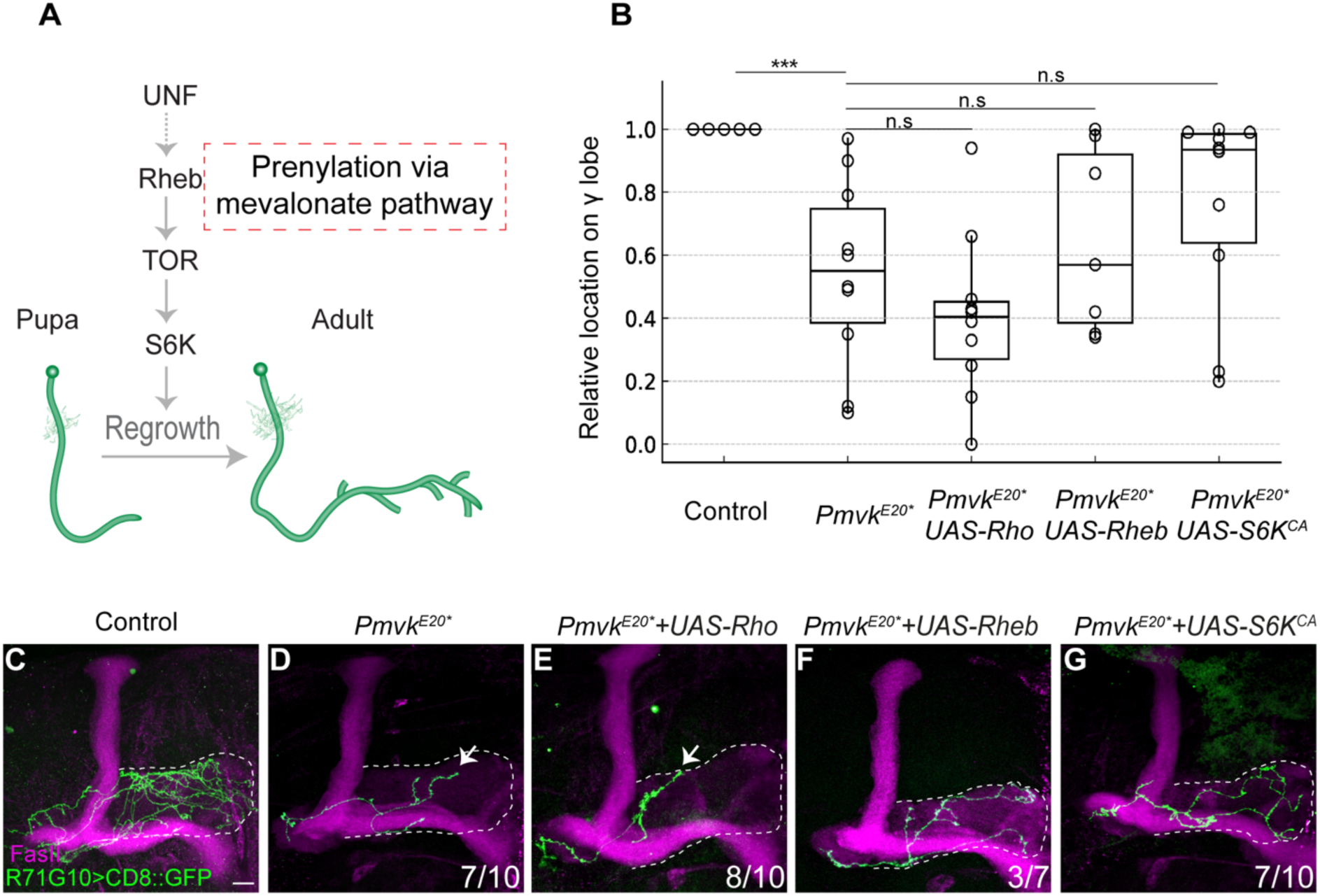
**Pmvk likely regulates axon regrowth via the TOR pathway**. (A) Scheme of developing γ-KCs in which UNF promotes regrowth via the TOR pathway (adapted from Yaniv et al., 2012). Rheb is a CAAX protein that undergoes prenylation, enabling its translocation to the plasma membrane to activate downstream effectors. (B) Quantification of axon regrowth for the genotypes displayed in (C-G). The x/y ratio indicates relative axon position along the adult γ lobe (quantification described in Figure S7). Box plots represent the median (line), interquartile range (box), and spread within 1.5×IQR (whiskers), circles indicate individual data points. Groups were compared using one-way ANOVA on logit-transformed data followed by Dunnett’s post-hoc test. A significant difference was observed only between control and *Pmvk^E20*^* (***p < 0.001). Comparisons of *Pmvk^E20*^* only with *Pmvk^E20*^* and UAS-Rho (p = 0.696), or UAS-Rheb (p = 0.704), or *UAS-S6K^CA^* (p = 0.120) were not significant. (C-G) Confocal Z-projections of γ single-cell MARCM clones in adult MBs, either control (C, comprised of multiple single cell clones), *Pmvk^E20*^* homozygous mutant (D), or *Pmvk^E20*^* homozygous mutant also overexpressing Rho (E), Rheb (F) or *S6K^CA^* (G). The γ-specific driver R71G10-Gal4 drives expression of membranal GFP (mCD8::GFP; green). Magenta is FasII staining. The γ lobe is outlined by a dashed white line. Arrows indicate axon growth defects. Numbers represent the fraction of MBs displaying the presented phenotype. Scale bar is 10 μm

### Pmvk likely regulates axon regrowth via the TOR pathway

The mevalonate pathway generates a common isoprenoid product that can be directed into five major downstream pathways (Figure 5D). In mammals, one of these is cholesterol biosynthesis, which is absent in *Drosophila*; instead, flies must obtain cholesterol from their diet (Cesur et al., 2023). The other four branches are: (2) ubiquinone synthesis, (3) dolichol synthesis, and (4–5) protein prenylation, through the addition of either farnesyl (4) or geranylgeranyl (5) groups. Prenylation increases protein hydrophobicity, thereby regulating their association with cellular membranes and influencing their function.

To test which of these branches might contribute to axon regrowth, we knocked down the major enzymes required for each process. However, none of these manipulations produced a regrowth defect (Figure S5). This could be due to insufficient efficiency of RNAi KD, or because simultaneous KD of two or more enzymes is necessary to reveal an effect. Since we were unable to pinpoint a specific downstream pathway, we decided to focus on the prenylation branch, as it was previously shown that prenylated proteins, such as the Rho GTPase and Rheb (Ras homolog enriched in brain), are associated with axon growth (Ng and Luo, 2004; Hall and Lalli, 2010).

Importantly, previous work from our lab demonstrated that the nuclear receptor unfulfilled (UNF) promotes axon regrowth by activating a program mediated, at least in part, by the Rheb-TOR-S6 kinase pathway (Figure 6A; Yaniv et al., 2012). We thus conducted rescue experiments in Pmvk mutants by overexpressing Rho, Rheb, and most importantly - a constitutively active variant of the Rheb downstream effector S6 kinase (S6K^CA^). Overexpression of Rho within Pmvk mutant clones showed no suppressive effect on the regrowth defect (Fig. 6B,E). Rheb overexpression did not significantly rescue γ-axon regrowth in the Pmvk mutant background (3/7 clones; Fig. 6B,F), which is consistent with the requirement for prenylation to achieve full Rheb activity and TOR activation. In contrast, S6K^CA^ overexpression showed an obvious trend toward suppression of the regrowth defect (7/10 clones displaying near-complete γ-axon extension; Fig. 6B,G). However, this effect did not reach statistical significance with the current sample size, which we unfortunately were unable to further increase (see acknowledgements). Taken together, these results suggest that Pmvk promotes γ-axon regrowth, at least in part, through the Rheb–TOR pathway.

## Discussion

How axon growth is regulated across different cellular and developmental contexts remains largely unknown. In this study, we used the remodeling MB as a model to uncover the transcriptional profiles of two lineage-related neuron types - γ-and α/β-KCs - which undergo developmental axon regrowth and initial axon growth, respectively, in similar spatiotemporal conditions. By comparing their transcriptional programs, we identified shared and unique expression patterns, highlighting candidates for an RNAi screen through which we identified genes promoting γ-and/or α/β-axon formation. One gene we found to be essential for regrowth of γ-axons is Pmvk, an enzyme in the mevalonate pathway responsible for isoprenoid biosynthesis. While Pmvk has not previously been directly linked to axon growth, isoprenoids are known to be required for the prenylation of small GTPases such as RhoA, Ras, and Cdc42—key regulators of the cytoskeleton and axon guidance (Hakeda-Suzuki et al., 2002; Hall and Lalli, 2010; Lee et al., 2021; Ng and Luo, 2004). Our findings suggest that Pmvk contributes to axon regrowth, at least in part, by supporting the membrane localization of Rheb, a prenylation-dependent GTPase previously shown to promote γ-axon regrowth (Yaniv et al., 2012).

A major strength of this study lies in its high temporal resolution, enabled by RNA-seq analysis across five developmental stages (at 3-hour intervals) plus adulthood. This was enabled due to a Gal4 line that we characterized and found to be specific to α/β-KCs throughout development, which allowed us to capture these neurons during active axon growth. Notably, our screening approach was aimed at uncovering shared genes required for both α/β growth and γ regrowth. However, despite identifying several γ regrowth-promoting genes, we did not uncover any strong candidates cell-autonomously required for α/β axon growth. This may reflect the robustness and/or redundancy of the genetic programs controlling α/β axon extension, where the loss of individual genes might be buffered by compensatory pathways. Previous studies suggested that larval γ axons provide a structural template for later-born KC types, including α/β-KCs, which follow a relatively stereotyped and stable developmental trajectory (Kurusu et al., 2002). Most of the genes known to affect α/β-KC development, such as FasII, Drl, and Netrin, impact guidance, branching, or stability but not necessarily initial axon extension (Bates et al., 2014; Fushima and Tsujimura, 2007; Hatch et al., 2021; Kang et al., 2019; Lin et al., 2009; Reynaud et al., 2015). One exception is twinstar (tsr), which encodes Cofilin and is required for early α/β axon growth, highlighting actin regulation as a potential neuron-intrinsic vulnerability in these cells (Ng and Luo, 2004). Similarly, Sickie was shown to be required for early α/β growth, functioning through a non-canonical Rac–Cofilin pathway (Abe et al., 2014). Notably, even in these cases, a scenario of normal initial growth followed by late developmental axonal degeneration has not been directly ruled out. Our findings suggest that combinatorial loss-of-function experiments, or temporally-targeted approaches, may be necessary to uncover additional regulators of α/β initial growth.

Our data provide a valuable resource for the *Drosophila* neuroscience community, particularly given that γ-and α/β-KCs together constitute the majority of the intrinsic MB neurons. The identification of Pmvk as a growth-promoting factor opens new avenues for dissecting how metabolic processes shape axonal remodeling. Since Pmvk is involved in isoprenoid synthesis required for protein prenylation, its role likely intersects with the activity of small GTPases such as Rheb, which are essential for growth signaling. This suggests a model in which metabolic enzymes like Pmvk act upstream to facilitate the activation of growth-promoting pathways through post-translational modification of key signaling molecules. Understanding this link between metabolism and signaling may shed light on how neurons integrate internal metabolic states with external developmental cues to regulate growth.

The broader role of the mevalonate pathway in axon growth is complex and context-dependent. While some studies have shown that inhibiting this pathway impairs axon extension, others suggest it may promote regeneration under certain conditions (Li et al., 2016; Schulz et al., 2004). This may reflect the dual role of the pathway in both isoprenoid and cholesterol synthesis. Pharmacological inhibition of the pathway aimed at reducing cholesterol levels —as with the widely used statins, which directly inhibit Hmgcr, the rate-limiting enzyme of the mevalonate pathway—could thus potentially have unintended effects on neuronal growth. Moreover, this pathway is tightly regulated by nutrient-sensitive signals, including insulin and TOR signaling (Belgacem and Martin, 2007), which activate SREBP, a transcription factor controlling lipid metabolism regulators like Hmgcr. In *Drosophila*, SREBP is known to regulate lipid homeostasis; therefore, it would be interesting to examine whether SREBP is required for neuronal remodeling (Seegmiller et al., 2002). Together, our findings highlight a critical link between cellular metabolism, signaling, and axon growth, suggesting that a neuron’s ability to remodel relies on sustained metabolic support and coordination between growth and nutrient-sensing pathways.

Finally, understanding axon growth capacity is key for understanding what limits axon regeneration following injury. Our transcriptomic and genetic analyses identify candidate genes that may therefore be relevant to axon repair following injury. Future efforts to injure γ-axons and compare their injury-evoked transcriptional program with those of initial α/β growth and developmental γ regrowth could provide valuable insights into shared and regeneration-specific molecular pathways.

## Acknowledgements

We thank the Bloomington Drosophila Stock Center and the Vienna Drosophila Resource Center for providing fly stocks. Monoclonal antibodies were obtained from the Developmental Studies Hybridoma Bank, created under the auspices of the NICHD and maintained by the University of Iowa. We are grateful to D. Robbins and to the staff of the Crown Institute for Genomics, INCPM for performing the RNA-seq, and to E. Hagai for assistance with cell sorting. This work was supported by the Minerva Foundation (grant # 714145) with funding from the Federal German Ministry of Education and Research, the Samowitz Foundation Trust, the Estate of Gerald Alexander, and by the European Research Council (ERC) Advanced Grant #101054886, “NeuRemodelBehavior”. O.S. is the incumbent of the Prof. Erwin Netter Professorial Chair of Cell Biology.

On June 15, 2025, our laboratory was directly hit by an Iranian missile and was completely destroyed, which unfortunately currently limits our ability to increase sample size (Figure 6) and to provide the Pmvk mutant allele generated in this study (which we are now in the process of rebuilding).

## References

Abe, T., Yamazaki, D., Murakami, S., Hiroi, M., Nitta, Y., Maeyama, Y., Tabata, T., 2014. The NAV2 homolog Sickie regulates F-actin-mediated axonal growth in Drosophila mushroom body neurons via the non-canonical Rac-Cofilin pathway. Development 141, 4716–4728. 10.1242/dev.113308

Alyagor, I., Berkun, V., Keren-Shaul, H., Marmor-Kollet, N., David, E., Mayseless, O., Issman-Zecharya, N., Amit, I., Schuldiner, O., 2018. Combining Developmental and Perturbation-Seq Uncovers Transcriptional Modules Orchestrating Neuronal Remodeling. Developmental Cell 47, 38–52.e6. 10.1016/j.devcel.2018.09.013

Bates, K.E., Sung, C., Hilson, L., Robinow, S., 2014. unfulfilled Interacting Genes Display Branch-Specific Roles in the Development of Mushroom Body Axons in Drosophila melanogaster. G3 Genes|Genomes|Genetics 4, 693–706. 10.1534/g3.113.009829

Belgacem, Y.H., Martin, J.-R., 2007. Hmgcr in the Corpus Allatum Controls Sexual Dimorphism of Locomotor Activity and Body Size via the Insulin Pathway in Drosophila. PLOS ONE 2, 1–11. 10.1371/journal.pone.0000187

Belle, J.S.D., Heisenberg, M., 1994. Associative odor learning in Drosophila abolished by chemical ablation of mushroom bodies. Science. 10.1126/science.8303280

Bornstein, B., Meltzer, H., Adler, R., Alyagor, I., Berkun, V., Cummings, G., Reh, F., Keren-Shaul, H., David, E., Riemensperger, T., Schuldiner, O., 2021. Transneuronal Dpr12/DIP-δ interactions facilitate compartmentalized dopaminergic innervation of Drosophila mushroom body axons. The EMBO Journal 40, e105763. 10.15252/embj.2020105763

Bornstein, B., Zahavi, E.E., Gelley, S., Zoosman, M., Yaniv, S.P., Fuchs, O., Porat, Z., Perlson, E., Schuldiner, O., 2015. Developmental Axon Pruning Requires Destabilization of Cell Adhesion by JNK Signaling. Neuron 88, 926–940. 10.1016/j.neuron.2015.10.023

Bradke, F., 2022. Mechanisms of Axon Growth and Regeneration: Moving between Development and Disease. J. Neurosci. 42, 8393. 10.1523/JNEUROSCI.1131-22.2022

Cesur, M.F., Basile, A., Patil, K.R., Çakır, T., 2023. A new metabolic model of <em>Drosophila melanogaster</em> and the integrative analysis of Parkinson’s disease. Life Sci. Alliance 6, e202201695. 10.26508/lsa.202201695

Dobin, A., Davis, C.A., Schlesinger, F., Drenkow, J., Zaleski, C., Jha, S., Batut, P., Chaisson, M., Gingeras, T.R., 2013. STAR: ultrafast universal RNA-seq aligner. Bioinformatics 29, 15–21. 10.1093/bioinformatics/bts635

Edwards, P.A., Ericsson, J., 2025. STEROLS AND ISOPRENOIDS: SIGNALING MOLECULES DERIVED FROM THE CHOLESTEROL BIOSYNTHETIC PATHWAY.

Fushima, K., Tsujimura, H., 2007. Precise control of fasciclin II expression is required for adult mushroom body development in Drosophila. Development, Growth & Differentiation 49, 215–227. 10.1111/j.1440-169X.2007.00922.x

Hakeda-Suzuki, S., Ng, J., Tzu, J., Dietzl, G., Sun, Y., Harms, M., Nardine, T., Luo, L., Dickson, B.J., 2002. Rac function and regulation during Drosophila development. Nature 416, 438–442. 10.1038/416438a

Hall, A., Lalli, G., 2010. Rho and Ras GTPases in Axon Growth, Guidance, and Branching. Cold Spring Harbor Perspectives in Biology 2, a001818–a001818. 10.1101/cshperspect.a001818

Hatch, H.A., Belalcazar, H.M., Marshall, O.J., Secombe, J., 2021. A KDM5–Prospero transcriptional axis functions during early neurodevelopment to regulate mushroom body formation. eLife 10, e63886. 10.7554/eLife.63886

Hilton, B.J., Husch, A., Schaffran, B., Lin, T., Burnside, E.R., Dupraz, S., Schelski, M., Kim, J., Müller, J.A., Schoch, S., Imig, C., Brose, N., Bradke, F., 2022. An active vesicle priming machinery suppresses axon regeneration upon adult CNS injury. Neuron 110, 51–69.e7. 10.1016/j.neuron.2021.10.007

Hinson, D.D., Chambliss, K.L., Toth, M.J., Tanaka, R.D., Gibson, K.M., 1997. Post-translational regulation of mevalonate kinase by intermediates of the cholesterol and nonsterol isoprene biosynthetic pathways. Journal of Lipid Research 38, 2216–2223. 10.1016/S0022-2275(20)34935-X

Kang, H., Zhao, J., Jiang, X., Li, G., Huang, W., Cheng, H., Duan, R., 2019. Drosophila Netrin-B controls mushroom body axon extension and regulates courtship-associated learning and memory of a Drosophila fragile X syndrome model. Molecular Brain 12, 52. 10.1186/s13041-019-0472-1

Kurusu, M., Awasaki, T., Masuda-Nakagawa, L.M., Kawauchi, H., Ito, K., Furukubo-Tokunaga, K., 2002. Embryonic and larval development of the Drosophila mushroom bodies: concentric layer subdivisions and the role of fasciclin II. Development 129, 409–419. 10.1242/dev.129.2.409

Lee, S.J., Zdradzinski, M.D., Sahoo, P.K., Kar, A.N., Patel, P., Kawaguchi, R., Aguilar, B.J., Lantz, K.D., McCain, C.R., Coppola, G., Lu, Q., Twiss, J.L., 2021. Selective axonal translation of the mRNA isoform encoding prenylated Cdc42 supports axon growth. Journal of Cell Science 134, jcs251967. 10.1242/jcs.251967

Lee, T., Lee, A., Luo, L., 1999. Development of the Drosophila mushroom bodies: Sequential generation of three distinct types of neurons from a neuroblast. Development 126, 4065– 4076.

Lee, T., Luo, L., 1999. Mosaic analysis with a repressible neurotechnique cell marker for studies of gene function in neuronal morphogenesis. Neuron. 10.1016/S0896-6273(00)80701-1

Lee, T., Marticke, S., Sung, C., Robinow, S., Luo, L., 2000. Cell-autonomous requirement of the USP/EcR-B ecdysone receptor for mushroom body neuronal remodeling in Drosophila. Neuron 28, 807–818. 10.1016/S0896-6273(00)00155-0

Li, H., Kuwajima, T., Oakley, D., Nikulina, E., Hou, J., Yang, W.S., Lowry, E.R., Lamas, N.J., Amoroso, M.W., Croft, G.F., Hosur, R., Wichterle, H., Sebti, S., Filbin, M.T., Stockwell, B., Henderson, C.E., 2016. Protein Prenylation Constitutes an Endogenous Brake on Axonal Growth. Cell Reports 16, 545–558. 10.1016/j.celrep.2016.06.013

Lin, S., Huang, Y., Lee, T., 2009. Nuclear Receptor Unfulfilled Regulates Axonal Guidance and Cell Identity of Drosophila Mushroom Body Neurons. PLOS ONE 4, 1–12. 10.1371/journal.pone.0008392

Lindner, J., Dassa, B., Wigoda, N., Stelzer, G., Feldmesser, E., Prilusky, J., Leshkowitz, D., 2025. UTAP2: an enhanced user-friendly transcriptome and epigenome analysis pipeline. BMC Bioinformatics 26, 79. 10.1186/s12859-025-06090-8

Love, M.I., Huber, W., Anders, S., 2014. Moderated estimation of fold change and dispersion for RNA-seq data with DESeq2. Genome Biology 15, 550. 10.1186/s13059-014-0550-8

Marmor-Kollet, N., Schuldiner, O., 2016. Contrasting developmental axon regrowth and neurite sprouting of Drosophila mushroom body neurons reveals shared and unique molecular mechanisms. Developmental Neurobiology 76, 262–276. 10.1002/dneu.22312

Ng, J., Luo, L., 2004. Rho GTPases Regulate Axon Growth through Convergent and Divergent Signaling Pathways. Neuron 44, 779–793. 10.1016/j.neuron.2004.11.014

Picelli, S., Faridani, O.R., Björklund, Å.K., Winberg, G., Sagasser, S., Sandberg, R., 2014. Full-length RNA-seq from single cells using Smart-seq2. Nature Protocols 9, 171–181. 10.1038/nprot.2014.006

Rabinovich, D., Yaniv, S.P., Alyagor, I., Schuldiner, O., 2016. Nitric Oxide as a Switching Mechanism between Axon Degeneration and Regrowth during Developmental Remodeling. Cell 164, 170–182. 10.1016/j.cell.2015.11.047

Reynaud, E., Lahaye, L.L., Boulanger, A., Petrova, I.M., Marquilly, C., Flandre, A., Martianez, T., Privat, M., Noordermeer, J.N., Fradkin, L.G., Dura, J.-M., 2015. Guidance of Drosophila Mushroom Body Axons Depends upon DRL-Wnt Receptor Cleavage in the Brain Dorsomedial Lineage Precursors. Cell Reports 11, 1293–1304. 10.1016/j.celrep.2015.04.035

Rozo, J.A., Martínez-Gallego, I., Rodríguez-Moreno, A., 2024. Cajal, the neuronal theory and the idea of brain plasticity. Frontiers in Neuroanatomy 18. 10.3389/fnana.2024.1331666

Schuldiner, O., Yaron, A., 2015. Mechanisms of developmental neurite pruning. Cellular and Molecular Life Sciences 72, 101–119. 10.1007/s00018-014-1729-6

Schulz, J.G., Bösel, J., Stoeckel, M., Megow, D., Dirnagl, U., Endres, M., 2004. HMG-CoA reductase inhibition causes neurite loss by interfering with geranylgeranylpyrophosphate synthesis. Journal of Neurochemistry 89, 24–32. 10.1046/j.1471-4159.2003.02305.x

Seegmiller, A.C., Dobrosotskaya, I., Goldstein, J.L., Ho, Y.K., Brown, M.S., Rawson, R.B., 2002. The SREBP Pathway in Drosophila: Regulation by Palmitate, Not Sterols. Developmental Cell 2, 229–238. 10.1016/S1534-5807(01)00119-8

Sudarsanam, S., Yaniv, S., Meltzer, H., Schuldiner, O., 2020. Cofilin regulates axon growth and branching of Drosophila γ-neurons. Journal of Cell Science 133. 10.1242/jcs.232595

Sutherland, T.C., Geoffroy, C.G., 2020. The Influence of Neuron-Extrinsic Factors and Aging on Injury Progression and Axonal Repair in the Central Nervous System. Frontiers in Cell and Developmental Biology Volume 8–2020. 10.3389/fcell.2020.00190

Xie, Q., Brbic, M., Horns, F., Kolluru, S.S., Jones, R.C., Li, J., Reddy, A.R., Xie, A., Kohani, S., Li, Z., McLaughlin, C.N., Li, T., Xu, C., Vacek, D., Luginbuhl, D.J., Leskovec, J., Quake, S.R., Luo, L., Li, H., 2021. Temporal evolution of single-cell transcriptomes of *Drosophila* olfactory projection neurons. eLife 10, e63450. 10.7554/eLife.63450

Yaniv, S.P., Issman-Zecharya, N., Oren-Suissa, M., Podbilewicz, B., Schuldiner, O., 2012. Axon regrowth during development and regeneration following injury share molecular mechanisms. Current Biology 22, 1774–1782. 10.1016/j.cub.2012.07.044

Yaniv, S.P., Meltzer, H., Alyagor, I., Schuldiner, O., 2020. Developmental axon regrowth and primary neuron sprouting utilize distinct actin elongation factors. Journal of Cell Biology 219. 10.1083/jcb.201903181

Zheng, B., Tuszynski, M.H., 2023. Regulation of axonal regeneration after mammalian spinal cord injury. Nature Reviews Molecular Cell Biology 24, 396–413. 10.1038/s41580-022-00562-y

Zhou, Y., Zhou, B., Pache, L., Chang, M., Khodabakhshi, A.H., Tanaseichuk, O., Benner, C., Chanda, S.K., 2019. Metascape provides a biologist-oriented resource for the analysis of systems-level datasets. Nature Communications 10, 1523. 10.1038/s41467-019-09234-6

