## Supplemental Figures S1-S7 and Supplementary Tables S1-S3 for "Comparative developmental transcriptomics of *Drosophila* mushroom body neurons highlights the mevalonate pathway as a regulator of axon growth"

### Supplemental material

| | Staining outside of the MB | α/β staining | α'/β' staining | γ staining |
|---|---|---|---|---|
| GMR16B01 | ✗ | ✓ | ✗ | ✗ |
| MB185B | ✓ | ✗ | ✗ | ✗ |
| MB594B | ✓ | ✗ | ✗ | ✓ |
| MB008B | ✗ | ✗ | ✗ | ✗ |
| MB477B | ✓ | ✓ | ✓ | ✗ |
| MB3714B | ✗ | ✗ | ✗ | ✗ |
| 17D | ✗ | ✗ | ✗ | ✗ |
| MZ1486 | ✗ | ✗ | ✗ | ✗ |
| C789 | ✓ | ✓ | ✓ | ✗ |
| GMR32E06 | ✗ | ✗ | ✗ | ✗ |
| NP3052 | ✓ | ✗ | ✗ | ✓ |
| GMR18F09 | ✓ | ✗ | ✓ | ✓ |
| GMR11A22 | ✓ | ✗ | ✗ | ✗ |
| GMR28H05 | ✗ | ✓ | ✓ | ✗ |
| GMR85D07 | ✗ | ✗ | ✗ | ✗ |
| GMR39H01 | ✓ | ✓ | ✓ | ✗ |
| GMR44E04 | ✓ | ✓ | ✗ | ✗ |
| GMR24B08 | ✓ | ✓ | ✓ | ✗ |
| GMR19E12 | ✓ | ✓ | ✓ | ✗ |
| GMR30D10 | ✓ | ✓ | ✓ | ✗ |
| GMR27G10 | ✓ | ✓ | ✓ | ✗ |
| GMR87B01 | ✓ | ✗ | ✗ | ✗ |
| GMR15H10 | ✓ | ✓ | ✓ | ✗ |
| GMR10B06 | ✓ | ✓ | ✓ | ✗ |
| NP6649 | ✗ | ✓ | ✓ | ✗ |
| NP3061 | ✓ | ✗ | ✓ | ✗ |
| GMR55D04 | ✗ | ✗ | ✗ | ✗ |
| VT042007 | ✓ | ✓ | ✓ | ✗ |
| VT0632326 | ✗ | ✗ | ✓ | ✓ |
| VT057379 | ✗ | ✗ | ✓ | ✓ |

**Table S1. Summary of α/β Gal4 screen – related to Figure 1**. We examined the expression of various Gal4 drivers - all considered as α/β specific in the adult - in the *Drosophila* brain at 24h APF by looking at Gal4-driven membrane-bound GFP (CD8::GFP). We scored for expression in the different intrinsic MB neurons (i.e. the three types of KCs), as well as outside the MB. The line eventually chosen for the cell sorting - R16B01-Gal4 (labeled in red) - is the only one specific to the MB and, within the MB, to α/β-KCs. The lines presented above were generated by the Janelia FlyLight Project Team in collaboration with the laboratories of Gerald M. Rubin and Barry J. Dickson. The MB lines were reported by Aso et al (2014), the GMR lines by Jenett et al (2012), the VT lines by Tirian and Dickson (2017), and the remaining seven lines by Aso et al (2009).

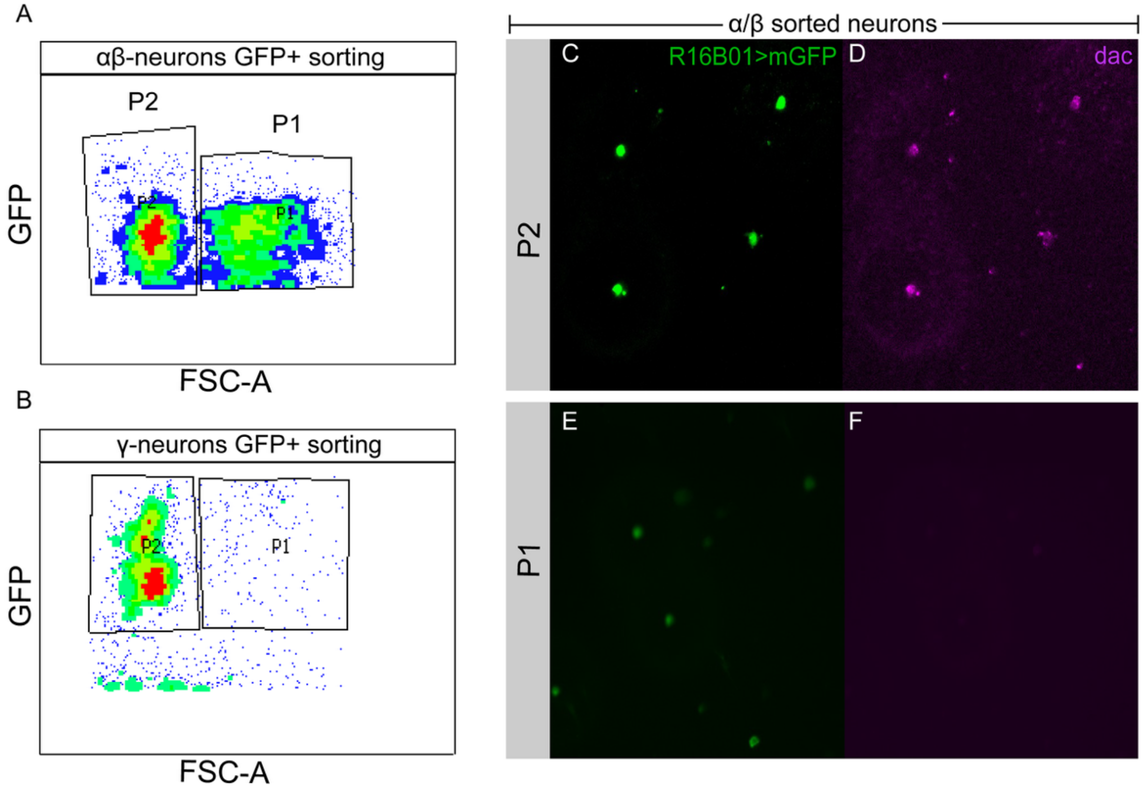


**Figure S1. Isolating α/β- and γ-KCs using fluorescence-activated cell sorter (FACS), related to Figure 1.** (A-B) FACS gating strategy for isolating GFP-positive α/β-KCs (A) or γ-KCs (B) from dissociated brains. Forward scatter (FSC) is plotted against GFP expression. P1 and P2 are different GFP-positive populations.

(C-F) Single confocal slices of the P2 (C-D) and P1 (E-F) post-sorted α/β-KCs labeled with mCD8-GFP driven by R16B01-Gal4 (green in C,E) and stained for Dachshund, a protein known to be expressed in all KCs (dac; magenta in D,F), direct us to conclude that P2 are α/β-KCs.

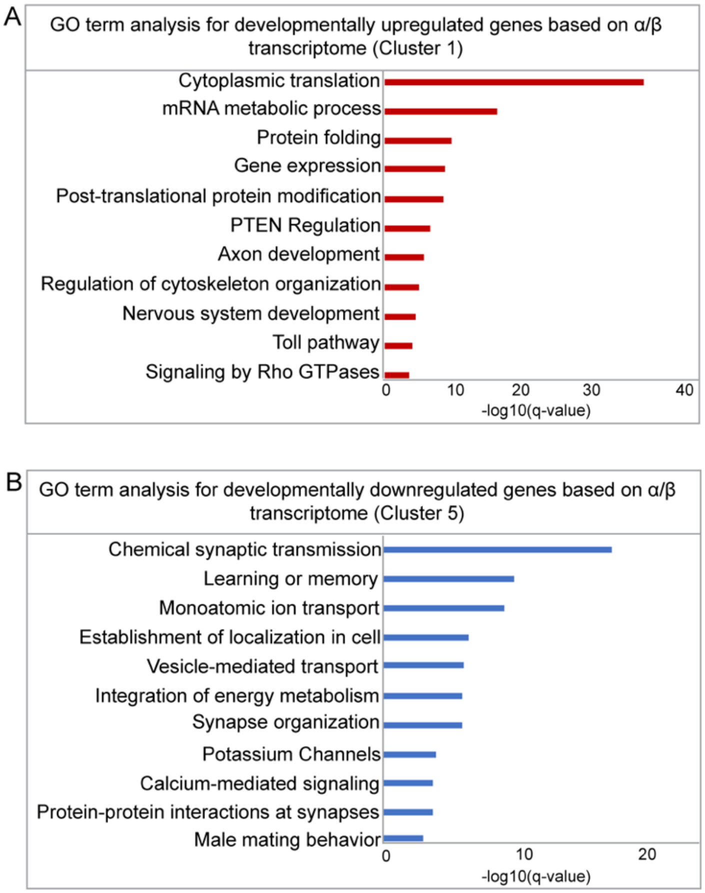


**Figure S2. Development-related terms are enriched in upregulated genes in the α/β transcriptome, related to Figure 2.**
(A-B) GO terms enriched in genes upregulated (A) or downregulated (B) during development, as identified from the α/β transcriptome (Clusters 1 or 5 in heatmap A, Figure 2, respectively).

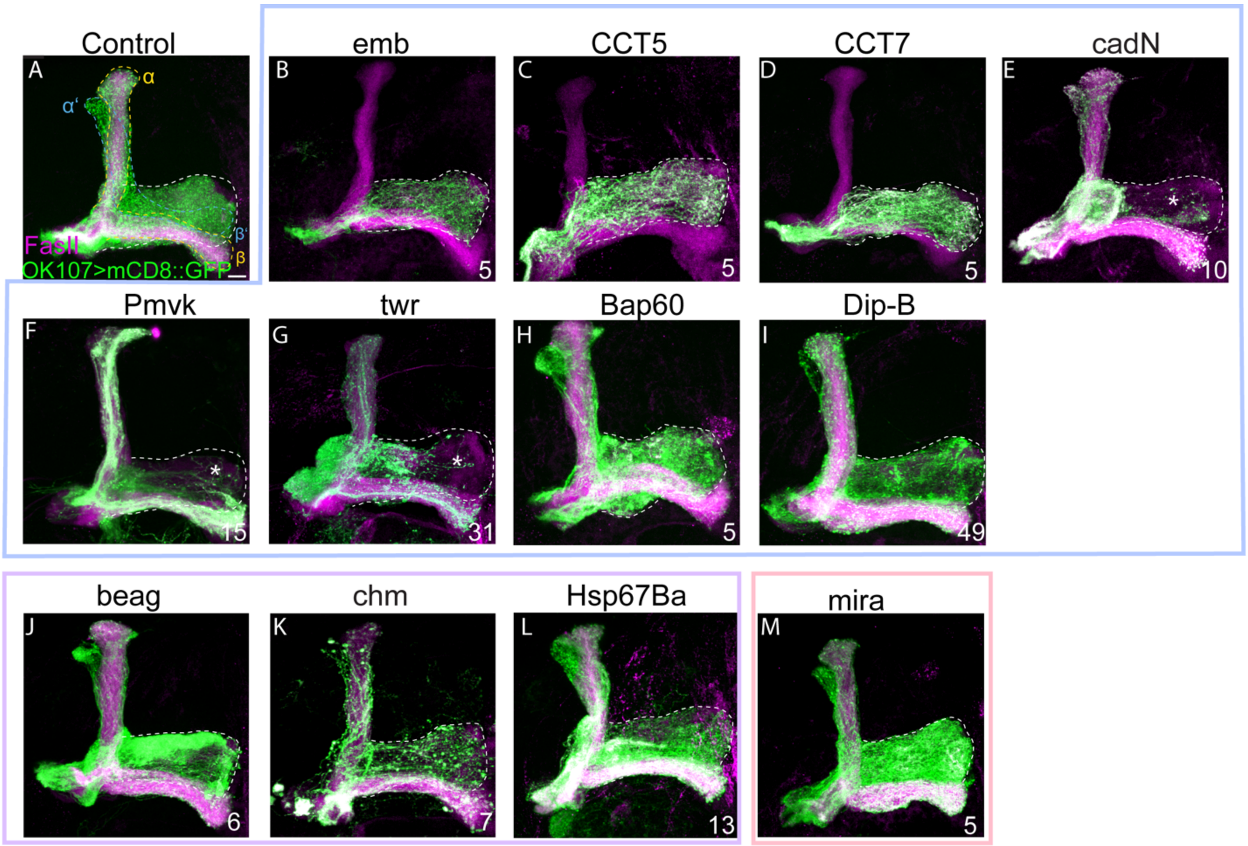


**Figure S3. Knocking down the 'positive hits' within MB clones revealed that CadN, Pmvk and twr are cell autonomously required for γ regrowth, related to Figure 3.** (A-M) Confocal Z-projections of adult KC neuroblast MARCM clones in control (A), or expressing RNAi targeting emb (B), CCT5 (C), CCT7 (D), cadN (E), Pmvk (F), twr (G), Bap60 (H), Dip-B (I), beag (J), chm(K) ,Hsp67Ba (L) , mira (M). OK107-Gal4 drives expression of the RNAi as well as membranal GFP (mCD8::GFP; green) in the clones. Magenta represents FasII staining. The full γ lobe is outlined in white and is based on the non-clonal γ-axons that are labeled with FasII. Asterisks mark the missing part of γ lobes. Frame colors in represent their phenotype as depicted in Figure 3R. Clones were generated by heat-shocking 24 hours after egg-laying. Numbers represent the sample size number. Scale bar is 10 μm.

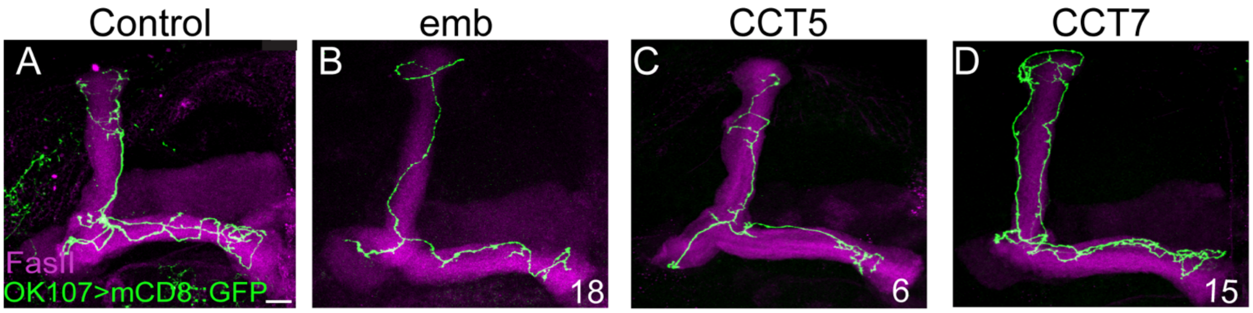


**Figure S4. KD of emb, CCT5, or CCT7 does not cell-autonomously affect initial α/β growth, related to Figure 3.** (A-D) Confocal Z-projections of adult α/β single-cell clones (or multiple single-cell clones) marked by membranal GFP (mCD8-GFP; green) driven by OK107-Gal4. Except for the control, OK017-Gal4 also drives expression of RNAi for the genes indicated above. Magenta is FasII staining. Clones were generated by heat shock at 0h APF. Numbers represent the sample size number. Scale bar is 10 μm.

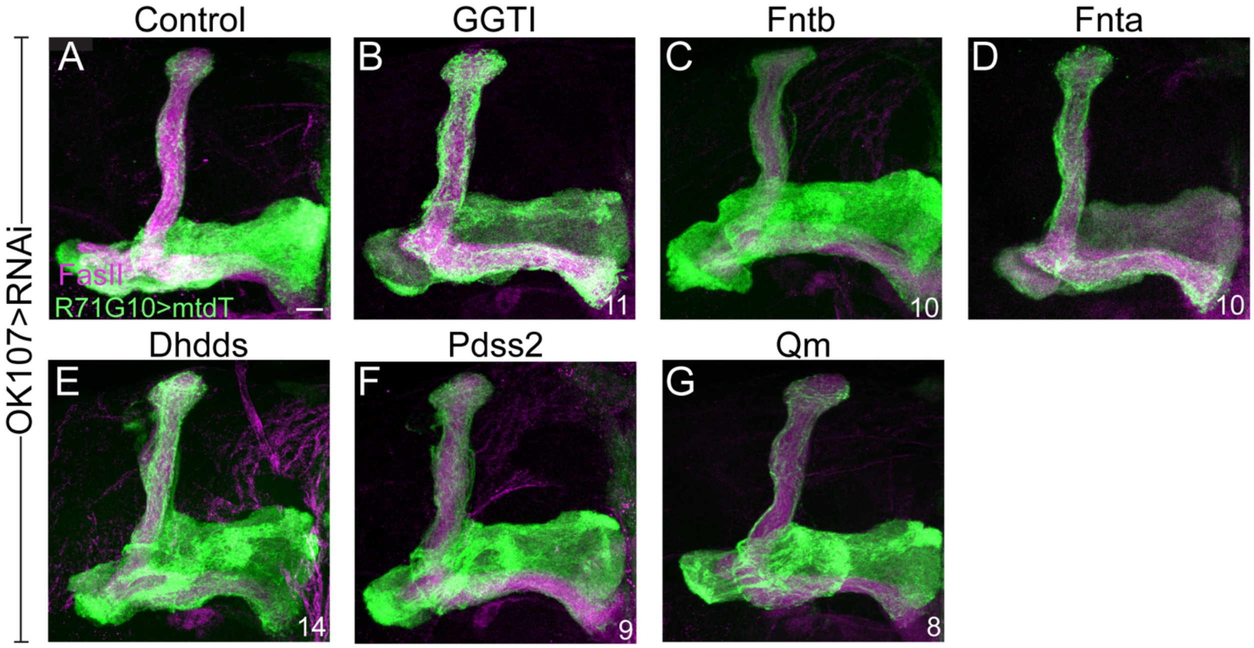


**Figure S5. KD of specific enzymes downstream of the mevalonate pathway does not affect regrowth of γ-axons, related to Figure 6**. (A-G) Confocal Z-projections of adult MBs of control (B) or those expressing RNAi targeting the indicated genes driven by OK107-Gal4. γ-KCs are labeled by mtdT:3xHA (green) driven by R71G10-QF2, while magenta indicates FasII staining. Numbers represent the sample size number. Scale bar is 10 µm.

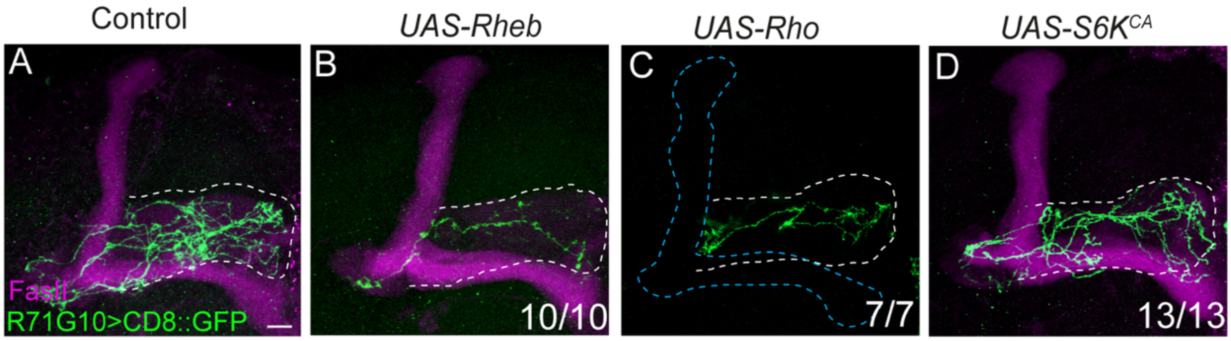


**Figure S6. Overexpression of Rheb, Rho or S6K$^{CA}$ does not affect axon regrowth, control panels for Figure 6.** (A-D) Confocal Z-projections of γ single-cell clones (or multiple single cell clones) in control brains (A), or those overexpressing *UAS-Rheb* (B), *UAS-Rho* (C), or *UAS-S6K$^{CA}$* (D) driven by the γ-specific driver R71G10-Gal4, which also drives expression of membranal GFP (mCD8::GFP; green). Magenta is FasII staining. The adult γ lobe is outlined by a dashed white line, as determined by FasII staining in A,B,D. In (C), FasII staining is absent due to a technical issue (see acknowledgments); therefore the outlines are of the expected γ lobe (white) and α/β lobes (blue). Numbers represent the fraction of MBs displaying the presented phenotype. Scale bar is 10 μm.

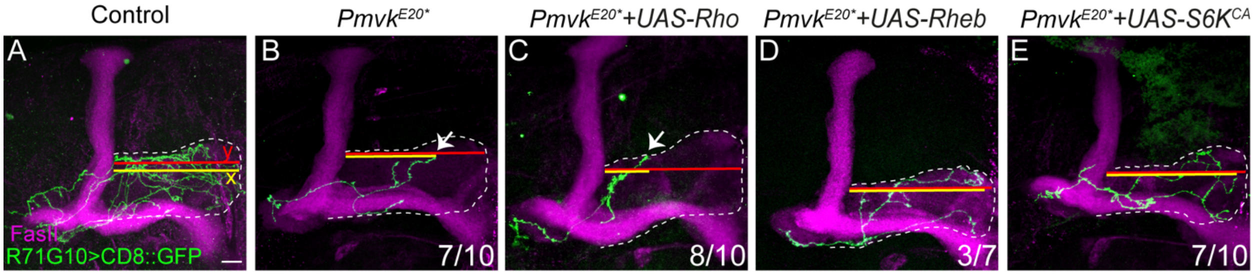


**Figure S7. Measurements of γ-axon regrowth, related to Fig 6**. (A-E) Confocal Z-projections of γ single-cell (or multiple single-cell) MARCM clones in adult MBs, either control (A), *Pmvk*$^{E20*}$ mutant (B), or *Pmvk*$^{E20*}$ mutant also overexpressing Rho (C), Rheb (D), or *S6K*$^{CA}$ (E). The γ-specific driver R71G10-Gal4 drives expression of membranal GFP (mCD8::GFP; green). Magenta is FasII staining. The γ lobe is outlined by a dashed white line. y (red) represents the length of the entire γ-lobe as determined by FasII staining; x (yellow) represents the extent of clonal γ-axon growth (the most distal point the axon reaches was chosen for quantification). Numbers represent the fraction of MBs displaying the presented phenotype. Scale bar is 10 μm.

| Gene name |
| --- |
| CG2915 |
| nito |
| Nmt |
| Cse1 |
| dl |
| Sar1 |
| Naa15-16 |
| Brd8 |
| Rack1 |
| CG42747 |
| Cyfip |
| Papss |
| CG17707 |
| HDAC1 |
| Hacd2 |
| Sumo |
| CG3815 |
| TER94 |
| Prp31 |
| CG3847 |
| Arpc2 |
| Gmd |
| Pep |
| me31B |
| jub |
| Mettl14 |
| MED15 |
| CG9801 |

**Table S2. Genes screened showing morphological defects but no axon growth, related to Figure 3.**

**Table S3. List of the 300 genes selected for the RNAi screen, related to Figure 3.**
All RNAi stocks were obtained from the Bloomington Drosophila Stock Center (BDSC).

| Gene name | RNAi Strain | Stock number |
|---|---|---|
| beta'COP | y[1] sc[*] v[1] sev[21]; P{y[+t7.7] v[+t1.8]=TRiP.HMS01038}attP2/TM3, Sb[1] | 36113 |
| beta'COP | y[1] v[1]; P{y[+t7.7] v[+t1.8]=TRiP.HM04017}attP2/TM3, Sb[1] | 31710 |
| CCT7 | y[1] sc[*] v[1] sev[21]; P{y[+t7.7] v[+t1.8]=TRiP.HMS01281}attP2 | 34931 |
| Mtpbeta | y[1] sc[*] v[1] sev[21]; P{y[+t7.7] v[+t1.8]=TRiP.HMS01017}attP2 | 34546 |
| Sec61alpha | y[1] sc[*] v[1] sev[21]; P{y[+t7.7] v[+t1.8]=TRiP.HMS01472}attP2 | 35730 |
| sel | y[1] v[1]; P{y[+t7.7] v[+t1.8]=TRiP.HMJ21136}attP40 | 51015 |
| Sema2b | y[1] v[1]; P{y[+t7.7] v[+t1.8]=TRiP.HM05143}attP2/TM3, Sb[1] | 28932 |
| Septin2 | y[1] v[1]; P{y[+t7.7] v[+t1.8]=TRiP.JF02838}attP2 | 28004 |
| Set | y[1] sc[*] v[1] sev[21]; P{y[+t7.7] v[+t1.8]=TRiP.HMC06565}attP40 | 77433 |
| SF1 | y[1] sc[*] v[1] sev[21]; P{y[+t7.7] v[+t1.8]=TRiP.HMC03680}attP40 | 52938 |
| Shc | y[1] sc[*] v[1] sev[21]; P{y[+t7.7] v[+t1.8]=TRiP.HMS05427}attP40 | 66961 |
| Slik | y[1] sc[*] v[1] sev[21]; P{y[+t7.7] v[+t1.8]=TRiP.HMC03770}attP40 | 55626 |
| SmF | y[1] v[1]; P{y[+t7.7] v[+t1.8]=TRiP.HMS05879}attP2/TM3, Sb[1] | 76067 |
| Smn | y[1] sc[*] v[1] sev[21]; P{y[+t7.7] v[+t1.8]=TRiP.HMS05688}attP2 | 67950 |
| Smn | y[1] sc[*] v[1] sev[21]; P{y[+t7.7] v[+t1.8]=TRiP.HMC03832}attP40 | 55158 |
| Snp | y[1] sc[*] v[1] sev[21]; P{y[+t7.7] v[+t1.8]=TRiP.HMS00321}attP2 | 33434 |
| Spn88Ea | y[1] v[1]; P{y[+t7.7] v[+t1.8]=TRiP.GL01344}attP40 | 42485 |
| Sps1 | y[1] v[1]; P{y[+t7.7] v[+t1.8]=TRiP.JF03230}attP2 | 29553 |
| Src64B | y[1] v[1]; P{y[+t7.7] v[+t1.8]=TRiP.HMC03327}attP40 | 51772 |
| Src64B | y[1] v[1]; P{y[+t7.7] v[+t1.8]=TRiP.JF03234}attP2 | 30517 |
| Ssrp | y[1] sc[*] v[1] sev[21]; P{y[+t7.7] v[+t1.8]=TRiP.HMS02285}attP2 | 41719 |
| Ssrp | y[1] v[1]; P{y[+t7.7] v[+t1.8]=TRiP.JF02120}attP2 | 26222 |
| ste24a | y[1] v[1]; P{y[+t7.7] v[+t1.8]=TRiP.HMC03454}attP40 | 51880 |
| Sumo | y[1] v[1]; P{y[+t7.7] v[+t1.8]=TRiP.JF02869}attP2 | 28034 |

| | | |
|---|---|---|
| Tapdelta | y[1] sc[*] v[1] sev[21]; P{y[+t7.7] v[+t1.8]=TRiP.HMC05581}attP40 | 64562 |
| Tbp | y[1] v[1]; P{y[+t7.7] v[+t1.8]=TRiP.GL00726}attP2 | 42770 |
| TER94 | y[1] v[1]; P{y[+t7.7] v[+t1.8]=TRiP.JF03402}attP2 | 31968 |
| TfIIB | y[1] sc[*] v[1] sev[21]; P{y[+t7.7] v[+t1.8]=TRiP.HMC05172}attP40 | 62165 |
| tw | y[1] sc[*] v[1] sev[21]; P{y[+t7.7] v[+t1.8]=TRiP.HMC03806}attP40 | 55735 |
| twe | y[1] sc[*] v[1] sev[21]; P{y[+t7.7] v[+t1.8]=TRiP.HMS00642}attP2 | 33044 |
| twe | y[1] sc[*] v[1] sev[21]; P{y[+t7.7] v[+t1.8]=TRiP.GL00547}attP2 | 36587 |
| twr | y[1] sc[*] v[1] sev[21]; P{y[+t7.7] v[+t1.8]=TRiP.HMS05732}attP40 | 67881 |
| Uch | y[1] v[1]; P{y[+t7.7] v[+t1.8]=TRiP.HMC06627}attP40 | 80391 |
| wuho | y[1] v[1]; P{y[+t7.7] v[+t1.8]=TRiP.HMJ23113}attP40 | 61281 |
| Wwox | y[1] v[1]; P{y[+t7.7] v[+t1.8]=TRiP.HMC02942}attP40 | 44546 |
| Wwox | y[1] v[1]; P{y[+t7.7] v[+t1.8]=TRiP.HMC03298}attP2 | 51747 |
| yki | y[1] v[1]; P{y[+t7.7] v[+t1.8]=TRiP.HMS00041}attP2 | 34067 |
| 14-3-3epsilon | y[1] v[1]; P{y[+t7.7] v[+t1.8]=TRiP.HM04001}attP2 | 31497 |
| ab | y[1] v[1]; P{y[+t7.7] v[+t1.8]=TRiP.JF03343}attP2 | 29407 |
| ago | y[1] sc[*] v[1] sev[21]; P{y[+t7.7] v[+t1.8]=TRiP.HMS00111}attP2 | 34802 |
| ago | y[1] v[1]; P{y[+t7.7] v[+t1.8]=TRiP.HM04005}attP2 | 31501 |
| Akap200 | y[1] v[1]; P{y[+t7.7] v[+t1.8]=TRiP.HM05018}attP2 | 28532 |
| AlkB | y[1] sc[*] v[1] sev[21]; P{y[+t7.7] v[+t1.8]=TRiP.HMS02673}attP40 | 43300 |
| aln | y[1] v[1]; P{y[+t7.7] v[+t1.8]=TRiP.JF03019}attP2 | 28382 |
| Arc42 | y[1] v[1]; P{y[+t7.7] v[+t1.8]=TRiP.HMC03503}attP40 | 53287 |
| Arp6 | y[1] sc[*] v[1] sev[21]; P{y[+t7.7] v[+t1.8]=TRiP.HMC05919}attP40 | 65155 |
| Arpc2 | y[1] v[1]; P{y[+t7.7] v[+t1.8]=TRiP.GL01469}attP2 | 43132 |
| Arpc2 | y[1] v[1]; P{y[+t7.7] v[+t1.8]=TRiP.HMJ23696}attP40/CyO | 62339 |
| Art1 | y[1] sc[*] v[1] sev[21]; P{y[+t7.7] v[+t1.8]=TRiP.GL01072}attP2/TM3, Sb[1] | 36891 |
| Art1 | y[1] v[1]; P{y[+t7.7] v[+t1.8]=TRiP.JF01306}attP2 | 31348 |
| aru | y[1] sc[*] v[1] sev[21]; P{y[+t7.7] v[+t1.8]=TRiP.HMS02966}attP2 | 50730 |
| aru | y[1] v[1]; P{y[+t7.7] v[+t1.8]=TRiP.HMJ21663}attP40 | 52976 |
| awd | y[1] v[1]; P{y[+t7.7] v[+t1.8]=TRiP.HMJ02099}attP40 | 42532 |
| Bap60 | y[1] v[1]; P{y[+t7.7] v[+t1.8]=TRiP.HMS00909}attP2 | 33954 |

| | | |
|---|---|---|
| beag | y[1] sc[*] v[1] sev[21]; P{y[+t7.7] v[+t1.8]=TRiP.HMC03533}attP2 | 53306 |
| bib | y[1] sc[*] v[1] sev[21]; P{y[+t7.7] v[+t1.8]=TRiP.HMC04807}attP2/TM3, Sb[1] | 57493 |
| bif | y[1] v[1]; P{y[+t7.7] v[+t1.8]=TRiP.JF03009}attP2 | 28372 |
| Bin1 | y[1] sc[*] v[1] sev[21]; P{y[+t7.7] v[+t1.8]=TRiP.HMC04115}attP40 | 56894 |
| Bin1 | y[1] sc[*] v[1] sev[21]; P{y[+t7.7] v[+t1.8]=TRiP.GL00127}attP2 | 36781 |
| brat | y[1] v[1]; P{y[+t7.7] v[+t1.8]=TRiP.HM05078}attP2 | 28590 |
| brat | y[1] v[1]; P{y[+t7.7] v[+t1.8]=TRiP.HM05078}attP2 | 28590 |
| Brd8 | y[1] sc[*] v[1] sev[21]; P{y[+t7.7] v[+t1.8]=TRiP.HMS02494}attP2 | 42658 |
| Brd8 | y[1] sc[*] v[1] sev[21]; P{y[+t7.7] v[+t1.8]=TRiP.HMC06412}attP40 | 67308 |
| Btk | y[1] v[1]; P{y[+t7.7] v[+t1.8]=TRiP.JF01797}attP2 | 25791 |
| CaBP1 | y[1] sc[*] v[1] sev[21]; P{y[+t7.7] v[+t1.8]=TRiP.HMC05746}attP40 | 64873 |
| Cad96Cb | y[1] v[1]; P{y[+t7.7] v[+t1.8]=TRiP.HMJ21472}attP40 | 54029 |
| Cad96Cb | y[1] v[1]; P{y[+t7.7] v[+t1.8]=TRiP.JF01881}attP2 | 25860 |
| CadN | y[1] sc[*] v[1] sev[21]; P{y[+t7.7] v[+t1.8]=TRiP.HMS02380}attP2 | 41982 |
| CCT2 | y[1] sc[*] v[1] sev[21]; P{y[+t7.7] v[+t1.8]=TRiP.HMS01190}attP2 | 34711 |
| CCT4 | y[1] sc[*] v[1] sev[21]; P{y[+t7.7] v[+t1.8]=TRiP.HMS05956}attP40/CyO | 77358 |
| CCT5 | y[1] v[1]; P{y[+t7.7] v[+t1.8]=TRiP.GL01246}attP2 | 41818 |
| CCT5 | y[1] sc[*] v[1] sev[21]; P{y[+t7.7] v[+t1.8]=TRiP.HMC04747}attP40 | 57440 |
| CG10038 | y[1] v[1]; P{y[+t7.7] v[+t1.8]=TRiP.HMJ21482}attP40 | 54039 |
| CG10465 | y[1] sc[*] v[1] sev[21]; P{y[+t7.7] v[+t1.8]=TRiP.HMC04553}attP40 | 57172 |
| CG10465 | y[1] v[1]; P{y[+t7.7] v[+t1.8]=TRiP.JF02026}attP2 | 26002 |
| CG10576 | y[1] sc[*] v[1] sev[21]; P{y[+t7.7] v[+t1.8]=TRiP.HMS01382}attP2 | 34388 |
| CG10602 | y[1] sc[*] v[1] sev[21]; P{y[+t7.7] v[+t1.8]=TRiP.GL00411}attP2/TM3, Sb[1] | 35482 |
| CG10602 | y[1] sc[*] v[1] sev[21]; P{y[+t7.7] v[+t1.8]=TRiP.HMC05668}attP40 | 64633 |
| CG10907 | y[1] sc[*] v[1] sev[21]; P{y[+t7.7] v[+t1.8]=TRiP.HMC03919}attP40 | 55204 |
| CG11134 | y[1] v[1]; P{y[+t7.7] v[+t1.8]=TRiP.JF03052}attP2 | 28637 |
| CG11360 | y[1] sc[*] v[1] sev[21]; P{y[+t7.7] v[+t1.8]=TRiP.HMC05165}attP40 | 62158 |

| | | |
|---|---|---|
| CG11360 | y[1] sc[*] v[1] sev[21]; P{y[+t7.7] v[+t1.8]=TRiP.HMS05762}attP40 | 67933 |
| CG11444 | y[1] sc[*] v[1] sev[21]; P{y[+t7.7] v[+t1.8]=TRiP.HMC06586}attP40 | 77454 |
| CG11964 | y[1] v[1]; P{y[+t7.7] v[+t1.8]=TRiP.HMJ30123}attP40 | 63557 |
| CG11999 | y[1] sc[*] v[1] sev[21]; P{y[+t7.7] v[+t1.8]=TRiP.HMS00565}attP2 | 34604 |
| CG12517 | y[1] v[1]; P{y[+t7.7] v[+t1.8]=TRiP.GLC01800}attP2 | 51742 |
| CG12708 | y[1] sc[*] v[1] sev[21]; P{y[+t7.7] v[+t1.8]=TRiP.HMC06559}attP40 | 77427 |
| CG13123 | y[1] v[1]; P{y[+t7.7] v[+t1.8]=TRiP.JF02423}attP2 | 29545 |
| CG13151 | y[1] v[1]; P{y[+t7.7] v[+t1.8]=TRiP.HMJ30001}attP40 | 62924 |
| CG13551 | y[1] sc[*] v[1] sev[21]; P{y[+t7.7] v[+t1.8]=TRiP.HMS01026}attP2 | 34554 |
| CG13667 | y[1] v[1]; P{y[+t7.7] v[+t1.8]=TRiP.HMJ24069}attP40/CyO | 62525 |
| CG14321 | y[1] v[1]; P{y[+t7.7] v[+t1.8]=TRiP.HMJ23812}attP40/CyO | 62389 |
| CG1440 | y[1] sc[*] v[1] sev[21]; P{y[+t7.7] v[+t1.8]=TRiP.HMC02912}attP2/TM3, Sb[1] | 44521 |
| CG14715 | y[1] v[1]; P{y[+t7.7] v[+t1.8]=TRiP.HMJ21596}attP40 | 52940 |
| CG1504 | y[1] sc[*] v[1] sev[21]; P{y[+t7.7] v[+t1.8]=TRiP.HMS05405}attP40 | 66939 |
| CG1529 | y[1] v[1]; P{y[+t7.7] v[+t1.8]=TRiP.GLC01412}attP40 | 44630 |
| CG1529 | y[1] sc[*] v[1] sev[21]; P{y[+t7.7] v[+t1.8]=TRiP.HMC05088}attP40 | 60094 |
| CG1674 | y[1] v[1]; P{y[+t7.7] v[+t1.8]=TRiP.HMJ23756}attP40/CyO | 62361 |
| CG17167 | y[1] v[1]; P{y[+t7.7] v[+t1.8]=TRiP.JF02560}attP2 | 27245 |
| CG17707 | y[1] v[1]; P{y[+t7.7] v[+t1.8]=TRiP.HMJ23942}attP40/CyO | 62461 |
| CG2852 | y[1] sc[*] v[1] sev[21]; P{y[+t7.7] v[+t1.8]=TRiP.HMC03892}attP40 | 55196 |
| CG2915 | y[1] v[1]; P{y[+t7.7] v[+t1.8]=TRiP.HMJ30194}attP40 | 63627 |
| CG3032 | y[1] sc[*] v[1] sev[21]; P{y[+t7.7] v[+t1.8]=TRiP.HMC04877}attP40 | 57560 |
| CG30379 | y[1] v[1]; P{y[+t7.7] v[+t1.8]=TRiP.JF02888}attP2 e[*] | 28988 |
| CG30379 | y[1] sc[*] v[1] sev[21]; P{y[+t7.7] v[+t1.8]=TRiP.HMS01567}attP2/TM3, Sb[1] | 36679 |
| CG30467 | y[1] v[1]; P{y[+t7.7] v[+t1.8]=TRiP.HMJ23664}attP40/CyO | 62307 |
| CG31548 | y[1] sc[*] v[1] sev[21]; P{y[+t7.7] v[+t1.8]=TRiP.HMC05773}attP40 | 64900 |
| CG31717 | y[1] sc[*] v[1] sev[21]; P{y[+t7.7] v[+t1.8]=TRiP.HMC06358}attP40 | 67255 |
| CG32187 | y[1] v[1]; P{y[+t7.7] v[+t1.8]=TRiP.HMJ30193}attP40/CyO | 63626 |
| CG3328 | y[1] sc[*] v[1] sev[21]; P{y[+t7.7] v[+t1.8]=TRiP.HMC04902}attP40 | 57713 |

| | | |
|---|---|---|
| CG3328 | y[1] v[1]; P{y[+t7.7] v[+t1.8]=TRiP.GLC01379}attP2 | 55211 |
| CG33303 | y[1] sc[*] v[1] sev[21]; P{y[+t7.7] v[+t1.8]=TRiP.HMC05782}attP40/CyO | 64909 |
| CG33721 | y[1] v[1]; P{y[+t7.7] v[+t1.8]=TRiP.HMJ23122}attP40 | 61290 |
| CG34353 | y[1] v[1]; P{y[+t7.7] v[+t1.8]=TRiP.HMJ22398}attP40 | 58291 |
| CG3815 | y[1] sc[*] v[1] sev[21]; P{y[+t7.7] v[+t1.8]=TRiP.HMS04327}attP40 | 56908 |
| CG3847 | y[1] sc[*] v[1] sev[21]; P{y[+t7.7] v[+t1.8]=TRiP.HMC04874}attP40 | 57557 |
| CG42346 | y[1] sc[*] v[1] sev[21]; P{y[+t7.7] v[+t1.8]=TRiP.HMC03255}attP2 | 51719 |
| CG42346 | y[1] v[1]; P{y[+t7.7] v[+t1.8]=TRiP.HM05169}attP2 | 28958 |
| CG42747 | y[1] sc[*] v[1] sev[21]; P{y[+t7.7] v[+t1.8]=TRiP.HMC06652}attP40 | 82974 |
| CG43658 | y[1] v[1]; P{y[+t7.7] v[+t1.8]=TRiP.JF03182}attP2 | 28754 |
| CG43658 | y[1] sc[*] v[1] sev[21]; P{y[+t7.7] v[+t1.8]=TRiP.HMS00332}attP2 | 32341 |
| CG4570 | y[1] sc[*] v[1] sev[21]; P{y[+t7.7] v[+t1.8]=TRiP.HMC04676}attP40 | 57378 |
| CG4709 | y[1] sc[*] v[1] sev[21]; P{y[+t7.7] v[+t1.8]=TRiP.HMC06318}attP40 | 67217 |
| CG5181 | y[1] sc[*] v[1] sev[21]; P{y[+t7.7] v[+t1.8]=TRiP.HMS00676}attP2 | 32888 |
| CG5721 | y[1] v[1]; P{y[+t7.7] v[+t1.8]=TRiP.HMJ21313}attP40 | 53949 |
| CG5880 | y[1] sc[*] v[1] sev[21]; P{y[+t7.7] v[+t1.8]=TRiP.HMC06290}attP2/TM3, Sb[1] | 66345 |
| CG5880 | y[1] sc[*] v[1] sev[21]; P{y[+t7.7] v[+t1.8]=TRiP.GL01121}attP2/TM3, Sb[1] | 36913 |
| CG5913 | y[1] v[1]; P{y[+t7.7] v[+t1.8]=TRiP.HMJ21347}attP40 | 53965 |
| CG6178 | y[1] v[1]; P{y[+t7.7] v[+t1.8]=TRiP.HM05128}attP2/TM3, Sb[1] | 28917 |
| CG6610 | y[1] sc[*] v[1] sev[21]; P{y[+t7.7] v[+t1.8]=TRiP.HM05263}attP2 | 31870 |
| CG7101 | y[1] sc[*] v[1] sev[21]; P{y[+t7.7] v[+t1.8]=TRiP.HMC06517}attP40 | 77386 |
| CG7369 | y[1] v[1]; P{y[+t7.7] v[+t1.8]=TRiP.GL01512}attP2 | 43170 |
| CG8031 | y[1] sc[*] v[1] sev[21]; P{y[+t7.7] v[+t1.8]=TRiP.HMC06643}attP40 | 80407 |
| CG8209 | y[1] v[1]; P{y[+t7.7] v[+t1.8]=TRiP.GL01249}attP2 | 41821 |
| CG8460 | y[1] v[1]; P{y[+t7.7] v[+t1.8]=TRiP.HMJ30020}attP40 | 62943 |
| CG8858 | y[1] sc[*] v[1] sev[21]; P{y[+t7.7] v[+t1.8]=TRiP.HMS01128}attP2 | 34975 |
| CG8860 | y[1] sc[*] v[1] sev[21]; P{y[+t7.7] v[+t1.8]=TRiP.HMC05121}attP40 | 60127 |

| | | |
|---|---|---|
| CG9135 | y[1] sc[*] v[1] sev[21]; P{y[+t7.7] v[+t1.8]=TRiP.HMS05617}attP40/CyO | 67856 |
| CG9588 | y[1] v[1]; P{y[+t7.7] v[+t1.8]=TRiP.HM05013}attP2 | 28527 |
| CG9667 | y[1] v[1]; P{y[+t7.7] v[+t1.8]=TRiP.HM04069}attP2 | 31758 |
| CG9801 | y[1] v[1]; P{y[+t7.7] v[+t1.8]=TRiP.HMS05308}attP40 | 63034 |
| CG9849 | y[1] sc[*] v[1] sev[21]; P{y[+t7.7] v[+t1.8]=TRiP.HMC04237}attP40 | 55948 |
| chm | y[1] sc[*] v[1] sev[21]; P{y[+t7.7] v[+t1.8]=TRiP.HMS00487}attP2 | 32484 |
| cib | y[1] v[1]; P{y[+t7.7] v[+t1.8]=TRiP.JF02837}attP2 | 28003 |
| ClC-b | y[1] v[1]; P{y[+t7.7] v[+t1.8]=TRiP.HMC03451}attP40 | 51877 |
| ClC-b | y[1] v[1]; P{y[+t7.7] v[+t1.8]=TRiP.JF01844}attP2 | 25826 |
| CNBP | y[1] sc[*] v[1] sev[21]; P{y[+t7.7] v[+t1.8]=TRiP.HMC03698}attP40 | 56971 |
| Cpsf6 | y[1] sc[*] v[1] sev[21]; P{y[+t7.7] v[+t1.8]=TRiP.HMS00113}attP2 | 34804 |
| Creld | y[1] sc[*] v[1] sev[21]; P{y[+t7.7] v[+t1.8]=TRiP.HMC06427}attP40 | 67323 |
| crim | y[1] sc[*] v[1] sev[21]; P{y[+t7.7] v[+t1.8]=TRiP.HMC06114}attP40 | 65362 |
| CRMP | y[1] v[1]; P{y[+t7.7] v[+t1.8]=TRiP.HMJ23963}attP40/CyO | 62479 |
| croc | y[1] sc[*] v[1] sev[21]; P{y[+t7.7] v[+t1.8]=TRiP.HMS01122}attP2 | 34647 |
| Cse1 | y[1] v[1]; P{y[+t7.7] v[+t1.8]=TRiP.JF02972}attP2 | 28337 |
| CSN4 | y[1] v[1]; P{y[+t7.7] v[+t1.8]=TRiP.GL01169}attP2 | 42798 |
| CSN5 | y[1] v[1]; P{y[+t7.7] v[+t1.8]=TRiP.GL01151}attP2 | 42781 |
| csul | y[1] sc[*] v[1] sev[21]; P{y[+t7.7] v[+t1.8]=TRiP.HMC04418}attP40 | 56978 |
| csul | y[1] v[1]; P{y[+t7.7] v[+t1.8]=TRiP.GL01544}attP40 | 43200 |
| Ctl1 | y[1] v[1]; P{y[+t7.7] v[+t1.8]=TRiP.HMJ23104}attP40 | 61272 |
| Cubn | y[1] v[1]; P{y[+t7.7] v[+t1.8]=TRiP.JF03118}attP2 | 28702 |
| Cubn | y[1] sc[*] v[1] sev[21]; P{y[+t7.7] v[+t1.8]=TRiP.HMS03167}attP2 | 51736 |
| Cyfip | y[1] sc[*] v[1] sev[21]; P{y[+t7.7] v[+t1.8]=TRiP.HMS01754}attP2 | 38294 |
| Cyp1 | y[1] sc[*] v[1] sev[21]; P{y[+t7.7] v[+t1.8]=TRiP.HMS00902}attP2 | 33950 |
| Cyp1 | y[1] v[1]; P{y[+t7.7] v[+t1.8]=TRiP.HMJ21213}attP40 | 53894 |
| D19B | y[1] v[1]; P{y[+t7.7] v[+t1.8]=TRiP.HMJ03125}attP40 | 51166 |
| Dad1 | y[1] sc[*] v[1] sev[21]; P{y[+t7.7] v[+t1.8]=TRiP.HMC05715}attP40 | 64680 |
| dap | y[1] sc[*] v[1] sev[21]; P{y[+t7.7] v[+t1.8]=TRiP.HMS01610}attP2 | 36720 |

| | | |
|---|---|---|
| Dci | y[1] sc[*] v[1] sev[21]; P{y[+t7.7] v[+t1.8]=TRiP.HMC05904}attP40 | 65030 |
| Ddx1 | y[1] v[1]; P{y[+t7.7] v[+t1.8]=TRiP.JF02682}attP2 | 27531 |
| dgt2 | y[1] v[1]; P{y[+t7.7] v[+t1.8]=TRiP.HM04038}attP2 | 31729 |
| dgt3 | y[1] v[1]; P{y[+t7.7] v[+t1.8]=TRiP.HMJ22086}attP40 | 58137 |
| Dip-B | y[1] v[1]; P{y[+t7.7] v[+t1.8]=TRiP.HMJ21488}attP40 | 54045 |
| dl | y[1] v[1]; P{y[+t7.7] v[+t1.8]=TRiP.JF02825}attP2 | 27650 |
| Dp1 | y[1] sc[*] v[1] sev[21]; P{y[+t7.7] v[+t1.8]=TRiP.HMS00659}attP2 | 32872 |
| drk | y[1] v[1]; P{y[+t7.7] v[+t1.8]=TRiP.JF02717}attP2 | 27563 |
| E(spl)malpha-BFM | y[1] sc[*] v[1] sev[21]; P{y[+t7.7] v[+t1.8]=TRiP.HMS05415}attP40 | 66949 |
| E(spl)mgamma-HLH | y[1] v[1]; P{y[+t7.7] v[+t1.8]=TRiP.JF02000}attP2 | 25978 |
| e(y)2 | y[1] v[1]; P{y[+t7.7] v[+t1.8]=TRiP.HMJ02090}attP40 | 42524 |
| eIF1 | y[1] sc[*] v[1] sev[21]; P{y[+t7.7] v[+t1.8]=TRiP.HMC04556}attP40 | 57174 |
| emb | y[1] sc[*] v[1] sev[21]; P{y[+t7.7] v[+t1.8]=TRiP.HMS00991}attP2 | 34021 |
| emb | y[1] v[1]; P{y[+t7.7] v[+t1.8]=TRiP.JF01311}attP2 | 31353 |
| ena | y[1] v[1]; P{y[+t7.7] v[+t1.8]=TRiP.JF01155}attP2 | 31582 |
| Eogt | y[1] sc[*] v[1] sev[21]; P{y[+t7.7] v[+t1.8]=TRiP.HMC06085}attP40 | 65224 |
| Ercc1 | y[1] v[1]; P{y[+t7.7] v[+t1.8]=TRiP.HMJ30048}attP40 | 62971 |
| Ercc1 | y[1] sc[*] v[1] sev[21]; P{y[+t7.7] v[+t1.8]=TRiP.GL01110}attP2 | 36906 |
| Ercc1 | y[1] v[1]; P{y[+t7.7] v[+t1.8]=TRiP.HMJ30048}attP40 | 62971 |
| ERp60 | y[1] sc[*] v[1] sev[21]; P{y[+t7.7] v[+t1.8]=TRiP.HMS00885}attP2 | 33935 |
| Ets65A | y[1] sc[*] v[1] sev[21]; P{y[+t7.7] v[+t1.8]=TRiP.HMS02246}attP2 | 41682 |
| Fadd | y[1] sc[*] v[1] sev[21]; P{y[+t7.7] v[+t1.8]=TRiP.HMS00123}attP2 | 34813 |
| Fdh | y[1] sc[*] v[1] sev[21]; P{y[+t7.7] v[+t1.8]=TRiP.HMS01268}attP2 | 34937 |
| FIG4 | y[1] sc[*] v[1] sev[21]; P{y[+t7.7] v[+t1.8]=TRiP.HMS01749}attP40 | 38291 |
| Flo1 | y[1] sc[*] v[1] sev[21]; P{y[+t7.7] v[+t1.8]=TRiP.GL00609}attP40 | 36649 |
| fpps | y[1] sc[*] v[1] sev[21]; P{y[+t7.7] v[+t1.8]=TRiP.HMC05747}attP40 | 64874 |
| fz3 | y[1] sc[*] v[1] sev[21]; P{y[+t7.7] v[+t1.8]=TRiP.GLC01626}attP2 | 44468 |

| | | |
|---|---|---|
| Galk | y[1] sc[*] v[1] sev[21]; P{y[+t7.7] v[+t1.8]=TRiP.GL00268}attP2 | 35356 |
| Galk | y[1] sc[*] v[1] sev[21]; P{y[+t7.7] v[+t1.8]=TRiP.HMC05085}attP40 | 60091 |
| Ge-1 | y[1] v[1]; P{y[+t7.7] v[+t1.8]=TRiP.GL00658}attP40/CyO | 38217 |
| Ge-1 | y[1] sc[*] v[1] sev[21]; P{y[+t7.7] v[+t1.8]=TRiP.HMS00340}attP2 | 32349 |
| Glg1 | y[1] sc[*] v[1] sev[21]; P{y[+t7.7] v[+t1.8]=TRiP.HMS01270}attP2 | 34921 |
| glo | y[1] sc[*] v[1] sev[21]; P{y[+t7.7] v[+t1.8]=TRiP.GL00438}attP40 | 36066 |
| glo | y[1] sc[*] v[1] sev[21]; P{y[+t7.7] v[+t1.8]=TRiP.HMS00079}attP2 | 33668 |
| Gmd | y[1] v[1]; P{y[+t7.7] v[+t1.8]=TRiP.HMJ21245}attP40 | 53912 |
| GMF | y[1] v[1]; P{y[+t7.7] v[+t1.8]=TRiP.HMC03184}attP40 | 51452 |
| gogo | y[1] sc[*] v[1] sev[21]; P{y[+t7.7] v[+t1.8]=TRiP.HMC05937}attP40 | 65193 |
| Gp210 | y[1] sc[*] v[1] sev[21]; P{y[+t7.7] v[+t1.8]=TRiP.HMC05931}attP40 | 65219 |
| Hacd2 | y[1] sc[*] v[1] sev[21]; P{y[+t7.7] v[+t1.8]=TRiP.HMS00109}attP2 | 34800 |
| HDAC1 | y[1] sc[*] v[1] sev[21]; P{y[+t7.7] v[+t1.8]=TRiP.HMS00607}attP2 | 33725 |
| HDAC1 | y[1] sc[*] v[1] sev[21]; P{y[+t7.7] v[+t1.8]=TRiP.HMS00164}attP2 | 34846 |
| Hel25E | y[1] sc[*] v[1] sev[21]; P{y[+t7.7] v[+t1.8]=TRiP.HMS00076}attP2 | 33666 |
| Hem | y[1] sc[*] v[1] sev[21]; P{y[+t7.7] v[+t1.8]=TRiP.HMS02252}attP2 | 41688 |
| HIP-R | y[1] sc[*] v[1] sev[21]; P{y[+t7.7] v[+t1.8]=TRiP.HMS00689}attP2 | 32900 |
| HIP-R | y[1] sc[*] v[1] sev[21]; P{y[+t7.7] v[+t1.8]=TRiP.HMS00988}attP2/TM3, Sb[1] | 34018 |
| His2Av | y[1] v[1]; P{y[+t7.7] v[+t1.8]=TRiP.HM05177}attP2 | 28966 |
| Hmgcr | y[1] v[1]; P{y[+t7.7] v[+t1.8]=TRiP.HMC03053}attP40 | 50652 |
| hpo | y[1] v[1]; P{y[+t7.7] v[+t1.8]=TRiP.HMS00006}attP2 | 33614 |
| hpo | y[1] v[1]; P{y[+t7.7] v[+t1.8]=TRiP.JF02740}attP2 | 27661 |
| Hrb87F | y[1] v[1]; P{y[+t7.7] v[+t1.8]=TRiP.JF01250}attP2/TM3, Ser[1] | 31472 |
| Hsp67Ba | y[1] v[1]; P{y[+t7.7] v[+t1.8]=TRiP.HMJ21725}attP40 | 53007 |
| Hsp67Ba | y[1] sc[*] v[1] sev[21]; P{y[+t7.7] v[+t1.8]=TRiP.HMS02359}attP2 | 41962 |
| Hsp67Bc | y[1] sc[*] v[1] sev[21]; P{y[+t7.7] v[+t1.8]=TRiP.HMS02440}attP40 | 42607 |

| | | |
|---|---|---|
| Hsp67Bc | y[1] sc[*] v[1] sev[21]; P{y[+t7.7] v[+t1.8]=TRiP.GL00377}attP2/TM3, Sb[1] | 35452 |
| Idh | y[1] v[1]; P{y[+t7.7] v[+t1.8]=TRiP.HMS02273}attP40 | 41708 |
| Ilp4 | y[1] v[1]; P{y[+t7.7] v[+t1.8]=TRiP.JF01346}attP2 | 31377 |
| inc | y[1] v[1]; P{y[+t7.7] v[+t1.8]=TRiP.HMJ22477}attP40 | 58346 |
| IntS9 | y[1] sc[*] v[1] sev[21]; P{y[+t7.7] v[+t1.8]=TRiP.HMC06154}attP40 | 65892 |
| jar | y[1] v[1]; P{y[+t7.7] v[+t1.8]=TRiP.JF02901}attP2 | 28064 |
| jub | y[1] v[1]; P{y[+t7.7] v[+t1.8]=TRiP.HMS02335}attP40 | 41938 |
| jub | y[1] sc[*] v[1] sev[21]; P{y[+t7.7] v[+t1.8]=TRiP.HMS00714}attP2 | 32923 |
| kn | y[1] v[1]; P{y[+t7.7] v[+t1.8]=TRiP.JF02206}attP2 | 31916 |
| kuk | y[1] v[1]; P{y[+t7.7] v[+t1.8]=TRiP.HMJ22805}attP40 | 60449 |
| kuk | y[1] sc[*] v[1] sev[21]; P{y[+t7.7] v[+t1.8]=TRiP.GLV21093}attP2 | 37487 |
| l(2)gd1 | y[1] v[1]; P{y[+t7.7] v[+t1.8]=TRiP.JF02619}attP2 | 27311 |
| Lac | y[1] v[1]; P{y[+t7.7] v[+t1.8]=TRiP.GL00666}attP40 | 38895 |
| Lac | y[1] v[1]; P{y[+t7.7] v[+t1.8]=TRiP.HM05151}attP2 | 28940 |
| Lam | y[1] v[1]; P{y[+t7.7] v[+t1.8]=TRiP.JF01389}attP2 | 31605 |
| lig | y[1] v[1]; P{y[+t7.7] v[+t1.8]=TRiP.HMJ23346}attP40 | 61857 |
| lncRNA:CR33938 | y[1] v[1]; P{y[+t7.7] v[+t1.8]=TRiP.HM05147}attP2/TM3, Sb[1] | 28936 |
| lost | y[1] sc[*] v[1] sev[21]; P{y[+t7.7] v[+t1.8]=TRiP.GL01090}attP2 | 38931 |
| Lrr47 | y[1] v[1]; P{y[+t7.7] v[+t1.8]=TRiP.HM05170}attP2/TM3, Sb[1] | 28959 |
| LSm3 | y[1] sc[*] v[1] sev[21]; P{y[+t7.7] v[+t1.8]=TRiP.HMC04113}attP40 | 56892 |
| Lst8 | y[1] sc[*] v[1] sev[21]; P{y[+t7.7] v[+t1.8]=TRiP.HMS01350}attP2 | 34361 |
| Lst8 | y[1] v[1]; P{y[+t7.7] v[+t1.8]=TRiP.GLC01412}attP40 | 44630 |
| Mad1 | y[1] sc[*] v[1] sev[21]; P{y[+t7.7] v[+t1.8]=TRiP.HMC03671}attP40 | 52930 |
| Mad1 | y[1] sc[*] v[1] sev[21]; P{y[+t7.7] v[+t1.8]=TRiP.GLV21088}attP2 | 35723 |
| mago | y[1] sc[*] v[1] sev[21]; P{y[+t7.7] v[+t1.8]=TRiP.HMC03947}attP40 | 55260 |
| mago | y[1] v[1]; P{y[+t7.7] v[+t1.8]=TRiP.HM05142}attP2 | 28931 |
| Map60 | y[1] v[1]; P{y[+t7.7] v[+t1.8]=TRiP.HMJ22424}attP40 | 58317 |
| Map60 | y[1] sc[*] v[1] sev[21]; P{y[+t7.7] v[+t1.8]=TRiP.HMS00457}attP2 | 32458 |
| mats | y[1] v[1]; P{y[+t7.7] v[+t1.8]=TRiP.JF03246}attP2 | 29567 |

| | | |
|---|---|---|
| mbo | y[1] sc[*] v[1] sev[21]; P{y[+t7.7] v[+t1.8]=TRiP.HMC06099}attP40 | 77374 |
| Mcm5 | y[1] sc[*] v[1] sev[21]; P{y[+t7.7] v[+t1.8]=TRiP.GL01301}attP40 | 41871 |
| me31B | y[1] v[1]; P{y[+t7.7] v[+t1.8]=TRiP.GL00695}attP40/CyO | 38923 |
| MED15 | y[1] sc[*] v[1] sev[21]; P{y[+t7.7] v[+t1.8]=TRiP.HMS00522}attP2 | 32517 |
| MED17 | y[1] sc[*] v[1] sev[21]; P{y[+t7.7] v[+t1.8]=TRiP.HMS01141}attP2 | 34664 |
| MED7 | y[1] sc[*] v[1] sev[21]; P{y[+t7.7] v[+t1.8]=TRiP.GL00472}attP2 | 35626 |
| MESR6 | y[1] v[1]; P{y[+t7.7] v[+t1.8]=TRiP.HMJ23813}attP40/CyO | 62390 |
| Mettl14 | y[1] sc[*] v[1] sev[21]; P{y[+t7.7] v[+t1.8]=TRiP.HMC05566}attP40 | 64547 |
| miple2 | y[1] sc[*] v[1] sev[21]; P{y[+t7.7] v[+t1.8]=TRiP.HMC04081}attP40 | 55393 |
| miple2 | y[1] v[1]; P{y[+t7.7] v[+t1.8]=TRiP.HMJ23667}attP40/CyO | 62310 |
| mira | y[1] sc[*] v[1] sev[21]; P{y[+t7.7] v[+t1.8]=TRiP.HMS00347}attP2 | 32356 |
| mof | y[1] sc[*] v[1] sev[21]; P{y[+t7.7] v[+t1.8]=TRiP.GL01011}attP40 | 36870 |
| Mondo | y[1] v[1]; P{y[+t7.7] v[+t1.8]=TRiP.JF02400}attP2 | 27059 |
| mop | y[1] v[1]; P{y[+t7.7] v[+t1.8]=TRiP.HM05008}attP2 | 28522 |
| mop | y[1] sc[*] v[1] sev[21]; P{y[+t7.7] v[+t1.8]=TRiP.HMS00706}attP2 | 32916 |
| Mppe | y[1] sc[*] v[1] sev[21]; P{y[+t7.7] v[+t1.8]=TRiP.HMC04967}attP40 | 57773 |
| msl-1 | y[1] sc[*] v[1] sev[21]; P{y[+t7.7] v[+t1.8]=TRiP.GL00256}attP2 | 35345 |
| mth | y[1] sc[*] v[1] sev[21]; P{y[+t7.7] v[+t1.8]=TRiP.HMS05784}attP40 | 67829 |
| mth | y[1] sc[*] v[1] sev[21]; P{y[+t7.7] v[+t1.8]=TRiP.GL01055}attP2 | 36823 |
| Mvd | y[1] sc[*] v[1] sev[21]; P{y[+t7.7] v[+t1.8]=TRiP.HMC06356}attP40 | 67253 |
| Naa15-16 | y[1] sc[*] v[1] sev[21]; P{y[+t7.7] v[+t1.8]=TRiP.HMS01400}attP2 | 34990 |
| Naglu | y[1] v[1]; P{y[+t7.7] v[+t1.8]=TRiP.HMC03368}attP40 | 51808 |
| NaPi-T | y[1] sc[*] v[1] sev[21]; P{y[+t7.7] v[+t1.8]=TRiP.HMS00966}attP2/TM3, Sb[1] | 34003 |
| Nct | y[1] sc[*] v[1] sev[21]; P{y[+t7.7] v[+t1.8]=TRiP.HMC04812}attP40 | 57497 |
| Nct | y[1] v[1]; P{y[+t7.7] v[+t1.8]=TRiP.JF02648}attP2 | 27498 |

| | | |
|---|---|---|
| NdufAF4 | y[1] v[1]; P{y[+t7.7] v[+t1.8]=TRiP.HM05053}attP2/TM3, Sb[1] | 28567 |
| Nedd8 | y[1] sc[*] v[1] sev[21]; P{y[+t7.7] v[+t1.8]=TRiP.HMS00818}attP2/TM3, Sb[1] | 33881 |
| nerfin-1 | y[1] v[1]; P{y[+t7.7] v[+t1.8]=TRiP.JF02956}attP2 | 28324 |
| nito | y[1] v[1]; P{y[+t7.7] v[+t1.8]=TRiP.HMJ02081}attP40 | 56852 |
| nito | y[1] v[1]; P{y[+t7.7] v[+t1.8]=TRiP.GLC01740}attP40 | 56860 |
| Nmt | y[1] v[1]; P{y[+t7.7] v[+t1.8]=TRiP.HMJ23820}attP40/CyO | 62397 |
| Nrt | y[1] v[1]; P{y[+t7.7] v[+t1.8]=TRiP.JF03170}attP2 | 28742 |
| Nsun5 | y[1] sc[*] v[1] sev[21]; P{y[+t7.7] v[+t1.8]=TRiP.HMS00438}attP2/TM3, Sb[1] | 32440 |
| Nsun5 | y[1] sc[*] v[1] sev[21]; P{y[+t7.7] v[+t1.8]=TRiP.HMS00780}attP2 | 32980 |
| Nup93-1 | y[1] v[1]; P{y[+t7.7] v[+t1.8]=TRiP.JF01712}attP2/TM3, Sb[1] | 31196 |
| Nup93-1 | y[1] v[1]; P{y[+t7.7] v[+t1.8]=TRiP.JF01712}attP2/TM3, Sb[1] | 31196 |
| Nup93-1 | y[1] sc[*] v[1] sev[21]; P{y[+t7.7] v[+t1.8]=TRiP.HMS00850}attP2 | 33908 |
| Orc4 | y[1] sc[*] v[1] sev[21]; P{y[+t7.7] v[+t1.8]=TRiP.HMS00404}attP2 | 32409 |
| orion | y[1] v[1]; P{y[+t7.7] v[+t1.8]=TRiP.HMJ22456}attP40 | 58326 |
| Ost48 | y[1] sc[*] v[1] sev[21]; P{y[+t7.7] v[+t1.8]=TRiP.HMS01303}attP2 | 34628 |
| OstDelta | y[1] sc[*] v[1] sev[21]; P{y[+t7.7] v[+t1.8]=TRiP.HMS01092}attP2/TM3, Sb[1] | 33752 |
| otk | y[1] v[1]; P{y[+t7.7] v[+t1.8]=TRiP.JF01796}attP2 | 25790 |
| p24-1 | y[1] sc[*] v[1] sev[21]; P{y[+t7.7] v[+t1.8]=TRiP.HMC04970}attP40 | 57776 |
| Pabp2 | y[1] sc[*] v[1] sev[21]; P{y[+t7.7] v[+t1.8]=TRiP.HMS00553}attP2 | 34602 |
| Paf-AHalpha | y[1] sc[*] v[1] sev[21]; P{y[+t7.7] v[+t1.8]=TRiP.HMC06063}attP40 | 65188 |
| Papss | y[1] v[1]; P{y[+t7.7] v[+t1.8]=TRiP.HMJ22841}attP40 | 60471 |
| Papss | y[1] sc[*] v[1] sev[21]; P{y[+t7.7] v[+t1.8]=TRiP.GL00291}attP2 | 35377 |
| Pax | y[1] v[1]; P{y[+t7.7] v[+t1.8]=TRiP.JF03111}attP2 | 28695 |
| pck | y[1] sc[*] v[1] sev[21]; P{y[+t7.7] v[+t1.8]=TRiP.GL00642}attP40 | 38203 |
| Pdi | y[1] v[1]; P{y[+t7.7] v[+t1.8]=TRiP.JF02874}attP2/TM3, Sb[1] | 28039 |
| Pen | y[1] v[1]; P{y[+t7.7] v[+t1.8]=TRiP.JF02772}attP2 | 27692 |
| Pep | y[1] sc[*] v[1] sev[21]; P{y[+t7.7] v[+t1.8]=TRiP.HMS00738}attP2 | 32944 |
| Pgant6 | y[1] sc[*] v[1] sev[21]; P{y[+t7.7] v[+t1.8]=TRiP.HMS00463}attP2 | 32463 |

| | | |
|---|---|---|
| PIG-C | y[1] sc[*] v[1] sev[21]; P{y[+t7.7] v[+t1.8]=TRiP.HMC06127}attP40 | 65375 |
| PIG-O | y[1] sc[*] v[1] sev[21]; P{y[+t7.7] v[+t1.8]=TRiP.HMC06349}attP40 | 67247 |
| PIG-V | y[1] v[1]; P{y[+t7.7] v[+t1.8]=TRiP.HMC03223}attP40 | 51478 |
| Pld3 | y[1] v[1]; P{y[+t7.7] v[+t1.8]=TRiP.JF01595}attP2 | 31122 |
| Pmvk | y[1] v[1]; P{y[+t7.7] v[+t1.8]=TRiP.HMJ21057}attP40/CyO | 50955 |
| pod1 | y[1] v[1]; P{y[+t7.7] v[+t1.8]=TRiP.JF01774}attP2 | 31219 |
| pod1 | y[1] sc[*] v[1] sev[21]; P{y[+t7.7] v[+t1.8]=TRiP.HMS02270}attP2 | 41705 |
| polo | y[1] sc[*] v[1] sev[21]; P{y[+t7.7] v[+t1.8]=TRiP.HMS00530}attP2 | 33042 |
| pont | y[1] v[1]; P{y[+t7.7] v[+t1.8]=TRiP.HMJ21078}attP40 | 50972 |
| por | y[1] sc[*] v[1] sev[21]; P{y[+t7.7] v[+t1.8]=TRiP.HMC04684}attP40 | 57380 |
| PPP4R2r | y[1] sc[*] v[1] sev[21]; P{y[+t7.7] v[+t1.8]=TRiP.HMS02416}attP40 | 42015 |
| PPP4R2r | y[1] v[1]; P{y[+t7.7] v[+t1.8]=TRiP.JF02065}attP2 | 26296 |
| Prdm13 | y[1] sc[*] v[1] sev[21]; P{y[+t7.7] v[+t1.8]=TRiP.HMC06382}attP40 | 67279 |
| prg | y[1] sc[*] v[1] sev[21]; P{y[+t7.7] v[+t1.8]=TRiP.GL00591}attP2/TM3, Sb[1] | 36631 |
| Prosalpha5 | y[1] sc[*] v[1] sev[21]; P{y[+t7.7] v[+t1.8]=TRiP.HMS00095}attP2 | 34786 |
| Prosbeta1 | y[1] sc[*] v[1] sev[21]; P{y[+t7.7] v[+t1.8]=TRiP.HMS00139}attP2/TM3, Sb[1] | 34824 |
| Prosbeta3 | y[1] sc[*] v[1] sev[21]; P{y[+t7.7] v[+t1.8]=TRiP.HMS00187}attP2 | 34868 |
| Prosbeta6 | y[1] sc[*] v[1] sev[21]; P{y[+t7.7] v[+t1.8]=TRiP.HMS00110}attP2 | 34801 |
| Prp19 | y[1] sc[*] v[1] sev[21]; P{y[+t7.7] v[+t1.8]=TRiP.HMS00652}attP2 | 32865 |
| Prp31 | y[1] sc[*] v[1] sev[21]; P{y[+t7.7] v[+t1.8]=TRiP.HMC04052}attP40 | 55364 |
| Prp8 | y[1] sc[*] v[1] sev[21]; P{y[+t7.7] v[+t1.8]=TRiP.HMS01297}attP2 | 34622 |
| Rab6 | y[1] v[1]; P{y[+t7.7] v[+t1.8]=TRiP.JF02640}attP2 | 27490 |
| Rack1 | y[1] sc[*] v[1] sev[21]; P{y[+t7.7] v[+t1.8]=TRiP.HMS01173}attP2 | 34694 |
| Ref1 | y[1] sc[*] v[1] sev[21]; P{y[+t7.7] v[+t1.8]=TRiP.HMS01301}attP2/TM3, Sb[1] | 34626 |
| Rgl | y[1] v[1]; P{y[+t7.7] v[+t1.8]=TRiP.HM05149}attP2 | 28938 |
| RhoGEF2 | y[1] v[1]; P{y[+t7.7] v[+t1.8]=TRiP.JF01747}attP2 | 31239 |

| | | |
|---|---|---|
| rig | y[1] sc[*] v[1] sev[21]; P{y[+t7.7] v[+t1.8]=TRiP.HMS01162}attP2 | 34684 |
| Rpn6 | y[1] v[1]; P{y[+t7.7] v[+t1.8]=TRiP.JF03317}attP2 | 29385 |
| Rpn6 | y[1] v[1]; P{y[+t7.7] v[+t1.8]=TRiP.JF03317}attP2 | 29385 |
| Rrp1 | y[1] sc[*] v[1] sev[21]; P{y[+t7.7] v[+t1.8]=TRiP.GL00343}attP2 | 35420 |
| Rrp1 | y[1] v[1]; P{y[+t7.7] v[+t1.8]=TRiP.HMJ23762}attP40/CyO | 62367 |
| ru | y[1] v[1]; P{y[+t7.7] v[+t1.8]=TRiP.HMJ21921}attP40 | 58065 |
| rump | y[1] sc[*] v[1] sev[21]; P{y[+t7.7] v[+t1.8]=TRiP.HMS02501}attP40 | 42665 |
| Sar1 | y[1] sc[*] v[1] sev[21]; P{y[+t7.7] v[+t1.8]=TRiP.HMS00355}attP2/TM3, Sb[1] | 32364 |
| SC35 | y[1] sc[*] v[1] sev[21]; P{y[+t7.7] v[+t1.8]=TRiP.HMC06150}attP40 | 65888 |
| Sce | y[1] sc[*] v[1] sev[21]; P{y[+t7.7] v[+t1.8]=TRiP.GL00371}attP2/TM3, Sb[1] | 35446 |
| Sce | y[1] v[1]; P{y[+t7.7] v[+t1.8]=TRiP.JF01396}attP2 | 31612 |
| Setx | y[1] sc[*] v[1] sev[21]; P{y[+t7.7] v[+t1.8]=TRiP.HMS01161}attP2/TM3, Sb[1] | 34683 |
| Sf3b6 | y[1] sc[*] v[1] sev[21]; P{y[+t7.7] v[+t1.8]=TRiP.HMS02566}attP40 | 42873 |
| Sf3b6 | y[1] sc[*] v[1] sev[21]; P{y[+t7.7] v[+t1.8]=TRiP.HMC03944}attP40 | 55257 |
| Shmt | y[1] sc[*] v[1] sev[21]; P{y[+t7.7] v[+t1.8]=TRiP.HMC04929}attP40 | 57739 |
| shv | y[1] sc[*] v[1] sev[21]; P{y[+t7.7] v[+t1.8]=TRiP.HMS01649}attP40 | 37507 |
| shv | y[1] v[1]; P{y[+t7.7] v[+t1.8]=TRiP.HMJ21390}attP40 | 54797 |
| SMC6 | y[1] sc[*] v[1] sev[21]; P{y[+t7.7] v[+t1.8]=TRiP.GL00500}attP2 | 36081 |
| SmF | y[1] v[1]; P{y[+t7.7] v[+t1.8]=TRiP.JF02276}attP2 | 26734 |
| Spn88Ea | y[1] sc[*] v[1] sev[21]; P{y[+t7.7] v[+t1.8]=TRiP.HMS01371}attP2 | 34381 |
| Sps1 | y[1] sc[*] v[1] sev[21]; P{y[+t7.7] v[+t1.8]=TRiP.GL00002}attP2 | 35135 |
| stet | y[1] sc[*] v[1] sev[21]; P{y[+t7.7] v[+t1.8]=TRiP.GLC01864}attP2 | 57698 |
| Sting | y[1] v[1]; P{y[+t7.7] v[+t1.8]=TRiP.JF01138}attP2 | 31565 |
| Stip1 | y[1] sc[*] v[1] sev[21]; P{y[+t7.7] v[+t1.8]=TRiP.HMS00779}attP2 | 32979 |
| tap | y[1] sc[*] v[1] sev[21]; P{y[+t7.7] v[+t1.8]=TRiP.HMS01395}attP2 | 34985 |
| Tbca | y[1] v[1]; P{y[+t7.7] v[+t1.8]=TRiP.JF01146}attP2 | 31573 |
| Tbca | y[1] v[1]; P{y[+t7.7] v[+t1.8]=TRiP.HMJ21731}attP40 | 53677 |

| | | |
|---|---|---|
| Tkt | y[1] v[1]; P{y[+t7.7] v[+t1.8]=TRiP.HMJ22552}attP40 | 60371 |
| Tmc | y[1] v[1]; P{y[+t7.7] v[+t1.8]=TRiP.HMJ21094}attP40 | 50984 |
| Tmem214 | y[1] sc[*] v[1] sev[21]; P{y[+t7.7] v[+t1.8]=TRiP.HMS01091}attP2 | 33751 |
| Uch-L5 | y[1] sc[*] v[1] sev[21]; P{y[+t7.7] v[+t1.8]=TRiP.HMC05716}attP40 | 64681 |
| Uch-L5 | y[1] sc[*] v[1] sev[21]; P{y[+t7.7] v[+t1.8]=TRiP.GL00357}attP2 | 35433 |
| Ugalt | y[1] v[1];P{y[+t7.7]v[+t1.8]=TRiP.HMC02402}attP2/TM3,Sb[1] | 44488 |
| Usp39 | y[1] sc[*] v[1] sev[21]; P{y[+t7.7] v[+t1.8]=TRiP.HMC03779}attP40 | 55632 |
| Yp1 | y[1] sc[*] v[1] sev[21]; P{y[+t7.7] v[+t1.8]=TRiP.HMC06320}attP40 | 67219 |
| Zw10 | y[1] sc[*] v[1] sev[21]; P{y[+t7.7] v[+t1.8]=TRiP.HMS02479}attP2/TM3, Sb[1] | 42643 |

**Table S3. List of the 300 genes selected for the RNAi screen, related to Figure 3.** All RNAi stocks were obtained from the Bloomington Drosophila Stock Center (BDSC).
